# GATA transcription factors drive initial Xist upregulation after fertilization through direct activation of a distal enhancer element

**DOI:** 10.1101/2022.08.02.502458

**Authors:** Liat Ravid Lustig, Abhishek Sampath Kumar, Till Schwämmle, Ilona Dunkel, Gemma Noviello, Raha Weigert, Guido Pacini, René Buschow, Afrah Ghauri, Maximilian Stötzel, Lars Wittler, Alexander Meissner, Edda G. Schulz

**Affiliations:** Systems Epigenetics, Otto Warburg Laboratories, Max Planck Institute for Molecular Genetics, 14195 Berlin, Germany; Department of Genome Regulation, Max Planck Institute for Molecular Genetics, 14195 Berlin, Germany; Microscopy and Cryo-Electron Microscopy, Max Planck Institute for Molecular Genetics, 14195 Berlin, Germany; Transgenic Unit, Max Planck Institute for Molecular Genetics, 14195 Berlin, Germany

## Abstract

To ensure dosage compensation for X-linked genes between the sexes, one X chromosome is silenced during early embryonic development of female mammals. This process of X-chromosome inactivation (XCI) is initiated through upregulation of the RNA *Xist* from one X chromosome shortly after fertilization. *Xist* then mediates chromosome-wide gene silencing in *cis* and remains expressed in all cell types except the germ line and the pluripotent state, where XCI is reversed. The factors that drive *Xist* upregulation and thereby initiate XCI remain however unknown. We identify GATA transcription factors as potent Xist activators and demonstrate that they are essential for the activation of *Xist* in mice following fertilization. Through a pooled CRISPR activation screen we find that GATA1 can drive ectopic *Xist* expression in murine embryonic stem cells (mESCs). We demonstrate that all GATA factors can activate *Xist* directly via a GATA-responsive regulatory element (RE79) positioned 100 kb upstream of the *Xist* promoter. Additionally, GATA factors are essential for the induction of XCI in mouse preimplantation embryos, as simultaneous deletion of three members of the GATA family (GATA1/4/6) in mouse zygotes effectively prevents *Xist* upregulation. Thus, initiation of XCI and possibly its maintenance in distinct lineages of the preimplantation embryo is ensured by the combined activity of different GATA family members, and the absence of GATA factors in the pluripotent state likely contributes to X reactivation. We thus describe a form of regulation in which the combined action of numerous tissue-specific factors can achieve near-ubiquitous expression of a target gene.

## Introduction

In female mammals, one out of two X chromosomes is silenced in a process called X-chromosome inactivation (XCI) (1). The master regulator of XCI, the long non-coding RNA Xist, is thus nearly ubiquitously expressed across tissues (2,3). In mice, Xist is upregulated shortly after fertilization and expressed in all cells with the exception of the pluripotent state and the germ line (4–6). However, the mechanism by which Xist upregulation is initially induced and then maintained remains largely unclear.

In mice, Xist is upregulated from the paternal X chromosome shortly after fertilization, while remaining repressed at the maternal allele due to a genomic imprint (4,5,7). This imprinted form of XCI (iXCI) is maintained in the extraembryonic tissues, but reversed in the pluripotent cells of the preimplantation embryo through Xist downregulation and loss of the imprint (4,5,8). This allows the transition from imprinted to random XCI, where each cell will inactivate either the paternal or the maternal X chromosome. Random XCI (rXCI) is then initiated shortly after implantation and maintained in all somatic cells (4,9). Murine embryonic stem cells (mESCs) are a cell culture model for the pluripotent cells of the preimplantation embryo and are widely used to study XCI, since female lines carry two active X chromosomes and initiate rXCI upon differentiation (10–14).

Xist expression is controlled by a large genomic region, which contains a series of lncRNA loci, thought to repress (Tsix, Linx) or activate (Jpx, Ftx, Xert) Xist transcription mostly in cis (15,16). Large (210-460 kb) single-copy Xist-containing transgenes (tg53, tg80), encompassing ~100kb genomic sequence upstream of Xist, can recapitulate initial Xist upregulation after fertilization and maintenance in extraembryonic lineages, but not rXCI in somatic tissues (17,18). Thus, Xist appears to be controlled in part by unique regulatory elements in different cellular settings. While enhancers responsible for post-fertilization Xist upregulation are unknown, we recently identified the functional enhancer repertoire responsible for Xist upregulation during rXCI (Gjaltema et al., 2022). The majority of the identified elements was indeed located outside the tg53/tg80 transgenes.

The enhancers that control Xist at the onset of rXCI are bound by several transcription factors (TFs) associated with the post-implantation pluripotent state such as OTX2 and SMAD2/3, which probably drive Xist upregulation in that developmental context (16). Downregulation of Xist at the pluripotent state, before the onset of rXCI, has been attributed to the repressive action of pluripotency factors, such as NANOG, REX1 (ZFP42), OCT4 (POU5F1) and PRDM14 (8,19–24). Since REX1 is already present throughout preimplantation development, XCI initiation after fertilization requires de-repression of Xist through the E3 ubiquitin ligase RNF12 (RLIM), which targets REX1 for degradation (24–26). However, the activating mechanisms that underlie the early upregulation of Xist upon fertilization remain unknown.

To identify Xist regulators, we used a pooled CRISPR activation (CRISPRa) screen in mESCs and identified a member of the GATA TF family as potent Xist activator. We then show that all family members can drive ectopic Xist upregulation. We identify a distal enhancer element that mediates GATA-dependent Xist expression, which is bound by different GATA TFs in extraembryonic cell lines. Finally, we demonstrate that a simultaneous zygotic knock-out of Gata1, Gata4 and Gata6 largely precludes Xist upregulation following fertilization in vivo. The joint action of different GATA TFs thus drives post-fertilization Xist upregulation and their absence in the epiblast might contribute to X reactivation.

## Results

### Pooled CRISPR screen identifies new Xist regulators

To identify unknown Xist activators, we performed a pooled CRISPRa screen to identify genes that, when overexpressed, can induce ectopic *Xist* upregulation. The screen was performed in male mESCs with a Tsix deletion (E14-STN_ΔTsixP_) and were differentiated by LIF withdrawal to sensitize cells for *Xis*t upregulation (Suppl. Fig. 1A). E14-STN_ΔTsixP_ cells also carry the SunTag CRISPRa system under control of a doxycycline-inducible promoter (Fig. 1A), which can induce strong ectopic upregulation, when recruited to the transcription start site (TSS) of a gene (27,28). We designed and cloned a custom lentiviral sgRNA library (CRISPRaX), targeting the promoters of both protein-coding and non-coding genes on the X chromosome, as well as known Xist regulators as controls (Suppl. Fig. 1B+C). We focused on X-chromosomal factors since X dosage is known to play an important role in *Xist* regulation. After transduction with the CRISPRaX library, leading to the genomic integration of a single sgRNA per cell, differentiating cells were stained for *Xist* RNA using Flow-FISH and the 15% of cells with the highest signal (Xist+) were enriched via flow cytometry (Fig. 1A). Genomic DNA was isolated from the sorted and unsorted populations, and the genomically integrated sgRNA sequences were quantified by Illumina sequencing (Suppl. Fig. 1D-F). To identify Xist regulators, we compared the abundance of sgRNAs in the sorted (Xist+) to the unsorted population using the MAGeCK MLE tool (29,30) (Fig. 1B-C, Suppl. Fig. 1G, Suppl. Table 1). The screen identified several known Xist activators, *Xist* itself (Fig. 1B, yellow) and a series of known repressors (Fig. 1C, red), confirming the sensitivity of the screen (19–22,31–35).

**Figure 1:**
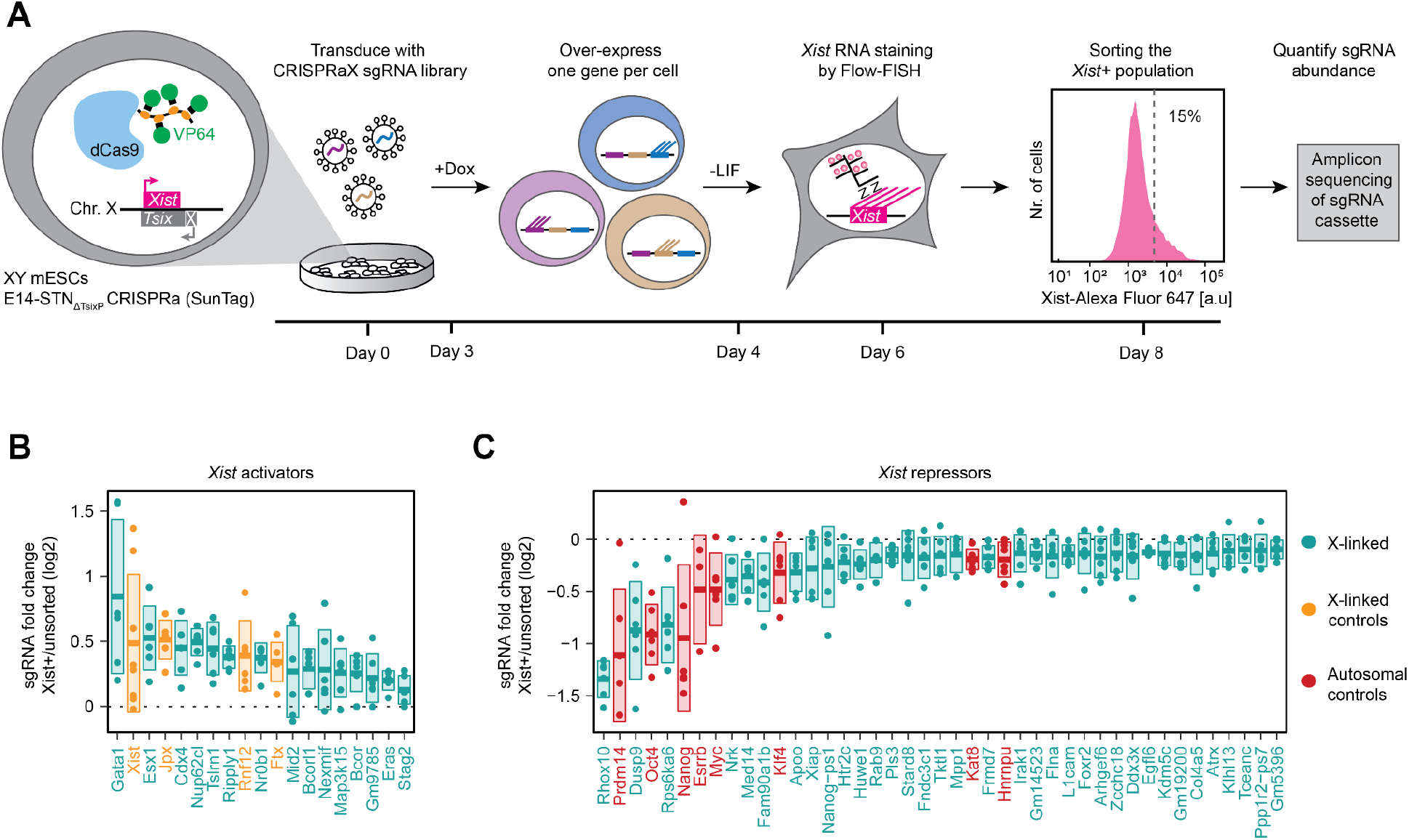
Pooled CRISPR activation screen identifies new Xist regulators. **(A)** Schematic depiction of the CRISPRa screen workflow. A male ESCl line with a deletion of the major *Tsix* promoter and a stably integrated doxycycline-inducible CRISPRa SunTag system (E14-STN_ΔTsix_) was transduced with a custom sgRNA library targeting X-chromosomal genes (CRISPRaX). Following puromycin selection, the cells were treated with doxycycline (Dox) to overexpress one gene per cell, and differentiated by LIF withdrawal (-LIF) to induce *Xist* upregulation. Cells were stained with Xist-specific probes by Flow-FISH and the top 15% Xist+ cells were sorted by flow cytometry. The sgRNA cassette was amplified from genomic DNA and sgRNA abundance in the unsorted and sorted populations was determined by deep sequencing. The screen was performed in three independent replicates. **(B-C)** Comparison of individual sgRNA abundance (dots) in the sorted fraction compared to the unsorted population for all significantly enriched (B) or depleted (C) genes in the screen (Wald-FDR<0.05, MAGeCK-MLE), colored as indicated. Genes are ordered by their beta-score, a measure for effect size (MAGeCK-MLE). The central line depicts the mean, boxes depict the standard deviation. Only the highest scoring TSS per gene is depicted.

### GATA1 is a potent Xist activator

Among the targeted X-linked genes, we found 15 putative activators, which were significantly enriched, and 35 putative repressors, which were depleted from the sorted fraction (Wald-FDR <0.05, MAGeCK, Fig. 1B-C, Suppl. Table 1). The top-scoring repressors were *Rhox10, Dusp9*, and *Rps6ka6* (Fig. 1C). While *Rhox10* has not yet been implicated in XCI to our knowledge, *Dusp9* and *Rps6ka6* likely interfere with *Xist* upregulation by delaying differentiation, as they are inhibitors of the differentiation-promoting MAPK signaling pathway (36–38). The top candidates as putative Xist activators were the transcription factors *Gata1, Cdx4*, the Rhox-family related gene *Esx1*, and the largely uncharacterized factor *Nup62cl* (Fig. 1B). To our knowledge, none of them has previously been implicated in *Xist* regulation or mESC differentiation. Only *Cdx4*, which lies ~150 kb downstream of the *Xist* gene, has been tested for a role in *Xist* regulation, but no effect could be detected upon deletion of its promoter (39). We validated the four top-scoring genes by individual overexpression, achieving >9-fold upregulation for all genes (Suppl. Fig. 2A+B).

While *Esx1, Cdx4* and *Nup62cl* overexpression only led to a small increase in Xist-expressing cells, *Gata1* induced strong *Xist* upregulation in the majority of cells (Suppl. Fig. 2C). Even in comparison to the positive control sgRNA, which activates the *Xist* promoter directly, *Gata1* overexpression resulted in more pronounced *Xist* upregulation. The Gata1-induced *Xist* distribution actually resembled the one seen in female mESCs upon differentiation (Suppl. Fig. 2D). Although *Xist* is thought to be repressed in undifferentiated mESCs, *Gata1* induced efficient *Xist* upregulation even without differentiation (Suppl. Fig. 2D). These observations suggest that GATA1 is an exceptionally strong Xist activator. We then inspected the expression dynamics of the identified putative activators during mESC differentiation within a previously generated RNA-seq data set (40). *Gata1, Cdx4* and *Esx1* showed very low or no expression at the time when *Xist* was upregulated (Suppl. Fig. 2E, Suppl. Table 2). Accordingly, knock-down of the strongest activator *Gata1* in female mESCs using CRISPR interference (CRISPRi) did not affect rXCI upon differentiation (Suppl. Fig. 2F-H). We therefore inspected expression of screen hits at other stages of early embryonic development, by re-analyzing published scRNA-seq data (41,42). *Gata1* was highly expressed between the 2-cell and the 16-cell stage (Suppl. Fig. 2I), suggesting a potential role in initial *Xist* upregulation upon fertilization.

### All GATA transcription factors are strong *Xist* activators

As GATA1 is part of a TF family with 6 members, which recognize similar DNA sequences (43), we tested whether the other family members might be able to induce *Xist* expression in a similar manner. To this end, we overexpressed all 6 GATA factors in male mESCs using CRISPRa (Fig. 2A), and measured their effect on *Xist* upregulation during differentiation. Each GATA factor could be overexpressed >150-fold, resulting in 35-65% Xist+ cells and 15-40-fold increase in *Xist* levels (Fig. 2B-F). Since some GATA factors have been shown to induce differentiation in mESCs (44,45), we tested whether they might activate *Xist* in an indirect manner through downregulation of pluripotency factors. We therefore assessed how GATA overexpression affected *Nanog, Oct4, Rex1, Esrrb* and *Prdm14* mRNA levels, but could not detect a consistent effect (Suppl Fig. 3A). GATA-mediated *Xist* induction can thus not be attributed to GATA-induced differentiation. We also tested, whether ectopic *Xist* upregulation upon GATA overexpression might be mediated by known Xist activators, but did again not observe any consistent effect on *Rnf12, Jpx, Ftx*, or *Yy1* (33–35,46) (Suppl. Fig. 3B). Since all GATA factors had a similar effect on *Xist*, we also analyzed whether they might induce each other’s expression. Here we indeed observed extensive cross-activation, where in particular *Gata4* and *Gata6* were induced by all other GATA factors (Suppl. Fig. 3C). Taken together, our results reveal that all six members of the GATA TF family are strong *Xist* activators, at least some of which might control *Xist* in a direct manner through activating the promoter or enhancer elements.

**Figure 2:**
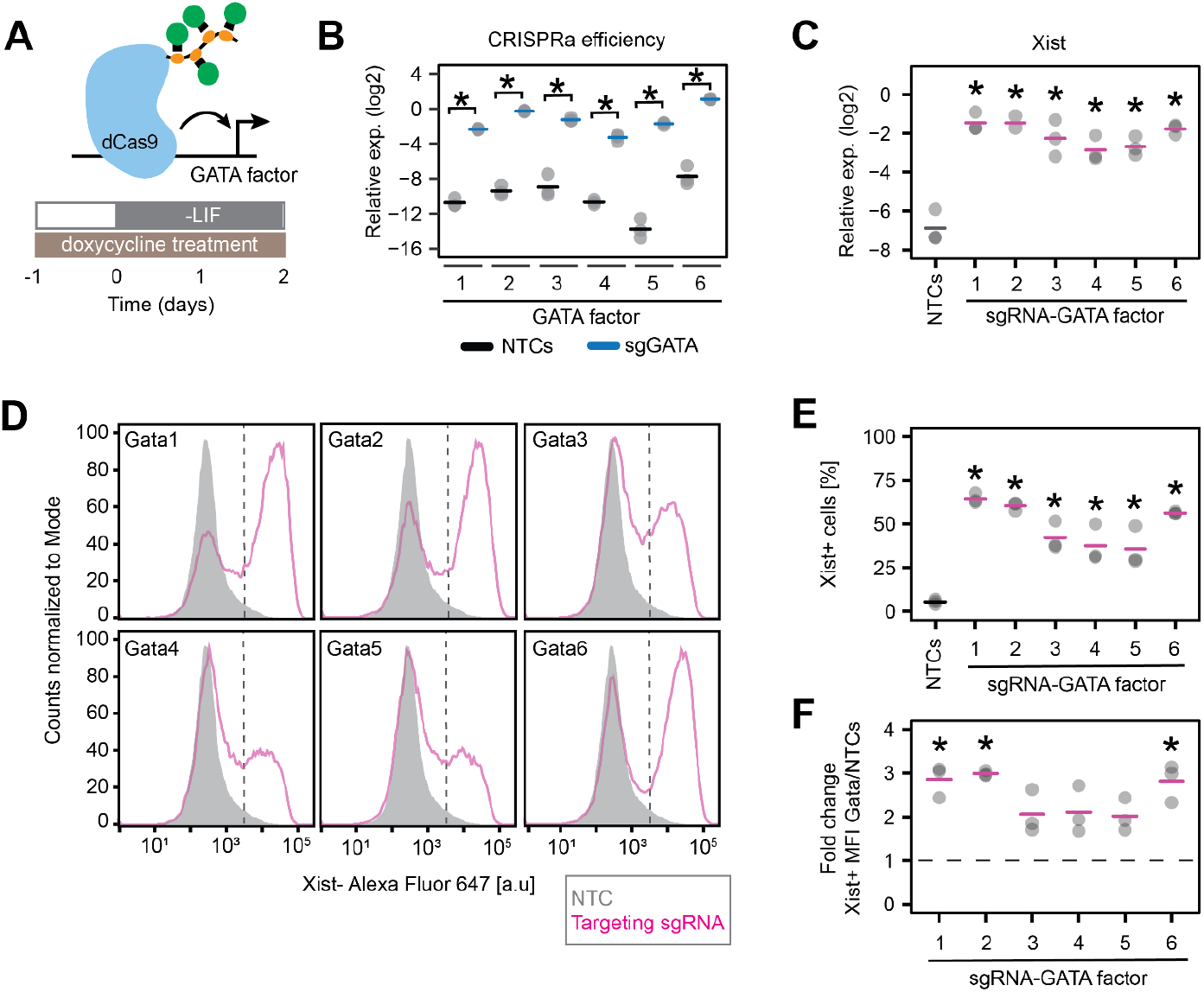
All GATA factors can induce *Xist* expression. **(A)** Schematic representation of the cell line (E14-STN_ΔTsixP_) and experimental setup used in B-F for ectopic overexpression of GATA family members. **(B-C)** Expression of GATA factors (B) and *Xist* (C) measured by qRT-PCR upon targeting each GATA TF by CRISPRa using 3 sgRNAs per gene. **(D-F)** Quantification of *Xist* by Flow-FISH. In (D) the sample shaded in grey denotes cells transduced with a non-targeting control (NTC) vector. Dashed lines divide Xist+ and Xist-cells, based on the 99th percentile of undifferentiated cells, transduced with NTCs, which do not express *Xist*. (F) Mean fluorescence intensity (MFI) within the Xist+ population of the targeted GATA factors compared to the NTC control. In (B,C,E,F) the mean (horizontal dashes) of 3 biological replicates (dots) is shown; asterisks indicate p<0.05 of a paired Student’s T-test for comparison to the respective NTC control.

### *Xist* is directly activated by GATA6 in a dose-dependent manner

To test whether a GATA TF could indeed directly induce *Xist* expression, we established a system that allowed rapid activation of a GATA family member to then follow the dynamics of *Xist* upregulation. We chose GATA6, because it is an important regulator of the primitive endoderm lineage, where *Xist* expression is maintained in an imprinted manner (47). We generated a female mESC line stably expressing HA-tagged *Gata6* cDNA N-terminally fused to the tamoxifen-inducible estrogen receptor (ERT2) domain (Fig. 3A). ERT2-GATA6 is retained in the cytoplasm and translocates into the nucleus upon treatment with 4-hydroxytamoxifen (4OHT) (Fig. 3B). The cells were cultured in 2i/LIF conditions, where *Xist* is repressed, and treated with 4OHT for 12 h. Starting at 6 h, we observed a significant increase in *Xist* levels (Fig. 3C), but no effect on the pluripotency factor *Nanog* (Suppl. Fig. 4A). We also assessed expression of three putative direct GATA6 target genes (48), two of which were significantly upregulated after 4 h of 4OHT treatment (*Sox7, Foxa2*, Suppl. Fig. 4B). The fact that upregulation of these genes only slightly precedes the upregulation of *Xist*, further supports the idea that GATA6 can directly induce *Xist*. We can however not exclude that other GATA6 target genes might additionally reinforce *Xist* upregulation.

**Figure 3:**
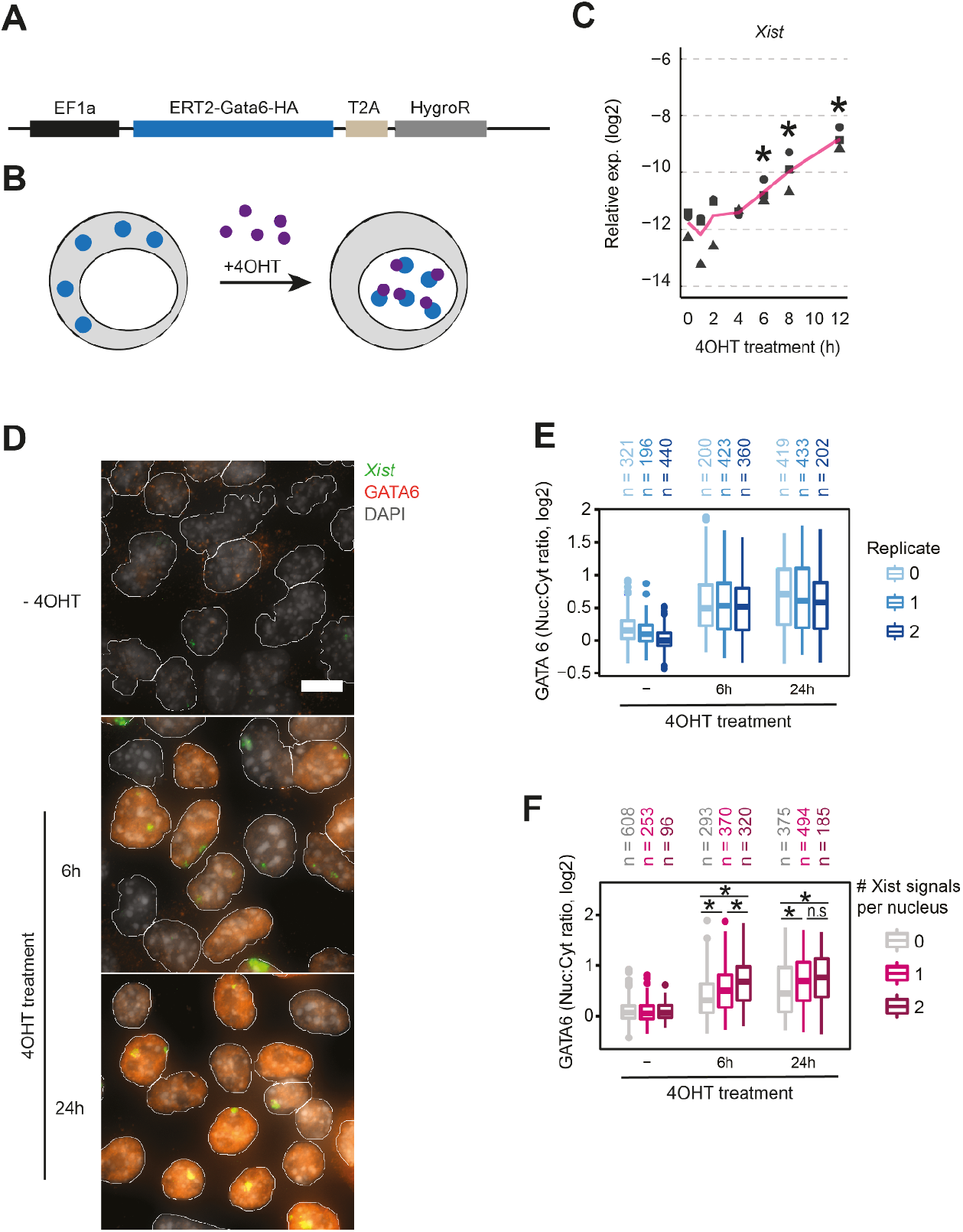
Xist is rapidly induced by GATA6 in a dose-dependent manner. **(A-B)** Schematic representation of the ERT2-GATA6 inducible system used in (C-F). Female TX-SP107 mESCs were transduced with a lentiviral vector shown in (A), expressing Gata6 cDNA N-terminally fused to the ERT2 domain and C-terminally tagged with HA. (B) Upon 4OHT treatment (purple), ERT2-GATA6-HA (blue) translocates into the nucleus. **(C)** Time course of Xist expression, assessed by qRT-PCR, upon 4OHT treatment of TX-SP107 ERT2-Gata6-HA cells, cultured in 2i/LIF medium. The line indicates the mean of 3 biological replicates (symbols); asterisks indicate p<0.05 using a paired Student’s T-test, comparing levels to the untreated control (Oh). **(D-F)** TX-SP107 ERT2-Gata6-HA cells, were grown on glass coverslips in conventional ESC medium (LIF only) for 48h and treated with 4OHT for 6 or 24h, followed by immunofluorescence staining (anti-HA to detect GATA6) combined with RNA-FISH (to detect Xist). In (E+F) nuclear translocation was quantified with automated image analysis. In (E) 3 biological replicates are shown, which were merged o for the analysis in (F) with excluding nuclei with >2 Xist signals. The central mark indicates the median, and the bottom and top edges of the box indicate the first and third quartiles, respectively. The top and bottom whiskers extend the boxes to a maximum of 1.5 times the interquartile range; cell numbers are indicated on top. In (F) asterisks indicate p<0.01, Wilcoxon ranksum test. Scale bar represents 10 μm.

To further characterize GATA6-dependent *Xist* regulation, we analyzed the relationship between nuclear GATA6 and *Xist* expression on the single cell level. To this end we performed immunofluorescence staining of HA-tagged ERT2-GATA6 combined with RNA-FISH for *Xist* (IF-FISH) after 6h and 24h of 4OHT treatment (Fig. 3D, Suppl. Fig. 4C). Through automated image segmentation, we quantified GATA6 staining within and around the nucleus to estimate nuclear and cytoplasmic staining intensities (see methods section for details). We then used the ratio between nuclear and cytoplasmic signals (nuc:cyt ratio) as a proxy for GATA6 accumulation in the nucleus. Although GATA6 expression levels appeared variable across cells, the nuc:cyt ratio was clearly increased in the majority of cells after 6h of 4OHT treatment, accompanied by an increase in *Xist*-expressing cells (Fig. 3E, Suppl. Fig. 4D). When analyzing the relationship between GATA6 levels and the *Xist* pattern, we observed that cells with higher GATA6 nuc:cyt ratios showed more *Xist* signals (Fig. 3F). These results show that GATA6 induces *Xist* in a dosage-dependent manner, further supporting direct regulation.

### GATA6 regulates *Xist* by binding to a distal enhancer element

Next, we aimed at identifying regulatory elements within *Xist*’s *cis*-regulatory landscape that mediate GATA-dependent regulation. As a first step, we identified binding sites for GATA factors in extraembryonic cell lines, which express different sets of GATA TFs and maintain *Xist* expression in an imprinted manner (10,11,49). We analyzed GATA2 and GATA3 in a female trophoblast stem (TS) cell line and GATA4 and GATA6 in an extraembryonic endoderm stem (XEN) cell line through CUT&Tag (50). In addition, we profiled the repressive histone modification H3K27me3, which has been shown to constitute the Xist imprint (7,51), and the active mark H3K27ac, which serves as a proxy for active enhancers (Fig. 4A, Suppl. Fig. 5, Suppl. Table 3).

**Figure 4:**
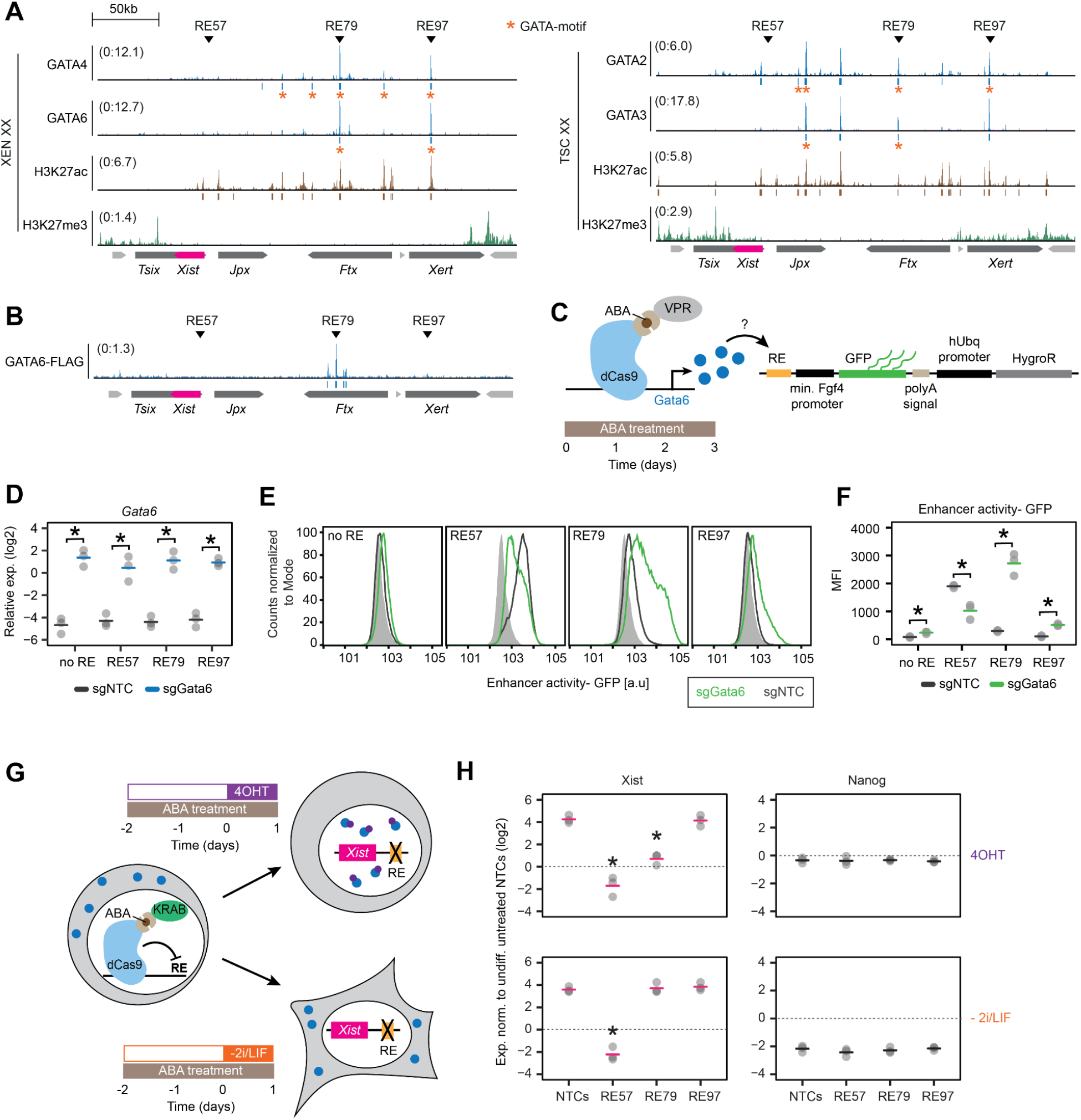
GATA6 regulates *Xist* by binding to a distal enhancer element. **(A)** Histone modifications and binding profiles for selected GATA TFs in female XEN (left) and TS cells (right), profiled by CUT&Tag. Peaks containing the respective GATA factor binding motif (p-value < 0.001, FIMO) are marked with an orange asterisk. 2-3 biological replicates were merged. **(B)** Published ChIP-seq data in mESCs overexpressing GATA6 (48). Arrowheads in (A+B), denote two regulatory elements (RE), RE79 and RE97, which are bound by all four tested GATA factors and the promoter-proxicmal RE57, which is not bound by GATA factors. Significant peaks (q-value < 0.05, MACS2) are indicated below the tracks. **(C-F)** Effect of GATA6 overexpression on a GFP reporter under control of different REs. TX-SP106 mESCs carrying a stably integrated ABA-inducible CRISPRa (VPR) system (C), were cultured in conventional ESC conditions and transduced with multiguide expression vectors of three sgRNAs against *Gata6* or with non-targeting controls (NTC). Cells were transduced with either the empty or RE-containing (RE57, RE79, RE97) lentiviral FIREWACh enhancer-reporter vector and treated with ABA for 3 days (C). Upregulation of *Gata6* was measured by qRT-PCR (D) and GFP levels were assessed by flow cytometry (E+F). In (E), light grey represents the cells’ autofluorescence. **(G-H)** Repression of REs through an ABA-inducible CRISPRi system and simultaneous GATA6 overexpression. Female TX-SP107 ERT2-Gata6-HA mESCs were cultured in 2i/LIF conditions and transduced with multiguide expression vectors of 3-4 sgRNAs against REs or with non-targeting controls (NTC). The cells were treated for 3 days with ABA to repress the respective RE and one day before harvesting, the cells were either differentiated (bottom, −2i/LIF, *Gata6*-independent *Xist* upregulation) or treated with 4OHT (top, *Gata6*-dependent *Xist* upregulation). *Xist* and *Nanog* mRNA levels were assessed by qRT-PCR. Samples were normalized to undifferentiated NTC controls not treated with 4OHT. In (D, F, H) horizontal dashes indicate the mean of 3 biological replicates (dots); asterisks indicate p<0.05 using a paired Student’s T-test for comparison to the respective NTC sample.

In both cell types we detected a series of H3K27ac peaks in a ~200kb region upstream of the *Xist* promoter, which was largely devoid of H3K27me3. Notably, this region has been shown to be covered by the maternal H3K27me3 imprint up to the blastocyst stage (7), further supporting the presence of *Xist* enhancers in that region. The maternal H3K27me3 domain however, appears to be lost in TS and XEN cells, in agreement with a previous study in TS cells (52). For the collected GATA binding profiles we performed a series of quality controls, since those factors, to our knowledge, have not been analyzed by CUT&Tag before (Suppl. Fig. 5, see methods for details). With the exception of GATA2, CUT&Tag appeared to primarily detect the expected binding sites. For GATA6 we observed two prominent binding sites in the 200 kb region upstream of *Xist*, both of which overlapped with H3K27ac peaks (Fig. 4A). These binding sites correspond to regulatory elements (RE) 79 and 97, which we have previously tested for *Xist* enhancer activity in differentiating mESCs through a pooled CRISPRi screen (16). RE97, but not RE79 was identified as a functional enhancer during the onset of rXCI in that screen. In a published GATA6 ChIP-seq data set, where GATA6 had been overexpressed in mESCs for 36 h, RE79 but not RE97 was strongly bound (Fig. 4B). However, both regions also appeared to be bound by GATA2 and GATA3 in TS cells and by GATA4 in XEN cells (Fig. 4A), supporting a functional role in GATA-mediated *Xist* expression.

To investigate whether GATA6 can indeed activate RE79 and RE97, we tested whether GATA6 overexpression could induce activation of a GFP reporter controlled by these potential enhancer elements. As a negative control, we also included RE57, which is located proximal to the *Xist* promoter and plays an important role in *Xist* regulation (16,53), but is not bound by GATA TFs (Fig. 4A). We cloned the three genomic regions (600-900bp) into a lentiviral enhancer-reporter plasmid, which was then co-expressed with a CRISPRa system to allow ectopic GATA6 upregulation (54,55) (Fig. 4C). A >30-fold overexpression of GATA6 indeed resulted in a strong 9- and 5-fold increase for RE79 and RE97, respectively, showing that these genomic loci can indeed act as GATA6-dependent enhancer elements (Fig. 4D-F). As expected, no increase in GFP levels upon GATA6 overexpression was detected for the RE57 reporter plasmid.

To test the functional importance of RE79 and RE97 in their endogenous genomic context, we next aimed to block their activation by CRISPRi and then probe the effect on GATA6-dependent *Xist* upregulation. To this end, we again made use of our female ERT2-GATA6 transgenic mESC line (Fig. 3) and co-expressed our CRISPRi system. Through simultaneous expression of 3-4 sgRNAs targeting one RE we blocked activation of RE79 and RE97 as well as the promoter-proximal RE57 as a control (Fig. 4G). Two days later, the cells were either treated with 4OHT to induce GATA6 translocation or differentiated to induce *Xist* upregulation in a GATA6-independent manner. Both, GATA6 induction (+4OHT) as well as differentiation (−2i/LIF) led to ~20-fold *Xist* upregulation in NTC-transduced control cells after 24 h (Fig. 4H). While targeting RE57 completely blocked *Xist* upregulation under both conditions, RE79 abolished GATA6-dependent *Xist* upregulation nearly completely (Fig. 4H, top), but did not affect differentiation-induced *Xist* expression (Fig. 4H, bottom). By contrast, targeting RE97 had no detectable effect in either context, suggesting that although RE97 can be bound and regulated by GATA factors, it does not regulate *Xist* via this mechanism in mESCs. The observation that RE97 also did not affect *Xist* expression upon 1 day of differentiation is in agreement with our previous finding that *Xist* is only affected by a deletion of the RE97-containing region from day 2 of differentiation onwards (16). Taken together, we can conclude that GATA6 induces *Xist* expression through binding to the RE79 element. Since this element is also bound by 3 other GATA TFs, we propose that it might mediate *Xist* regulation by all family members.

### GATA factors are required for *Xist* upregulation after fertilization *in vivo*

Having demonstrated the potency of GATA factors as Xist activators, we examined the physiological significance of GATA-dependent *Xist* regulation. To this end, we first analyzed GATA expression patterns during early development at the level of transcripts and proteins through re-analysis of published single-cell RNA-seq data (41,56) and immunofluorescence staining (Fig. 5A-B, Suppl. Fig. 6). In agreement with previous reports, multiple GATA factors were expressed at all stages of preimplantation development with exception of the pluripotent epiblast (47). This expression profile perfectly mirrors the reported pattern of imprinted *Xist* expression, which is upregulated shortly after fertilization, and downregulated only in pluripotent cells (4,5,57).

**Figure 5.**
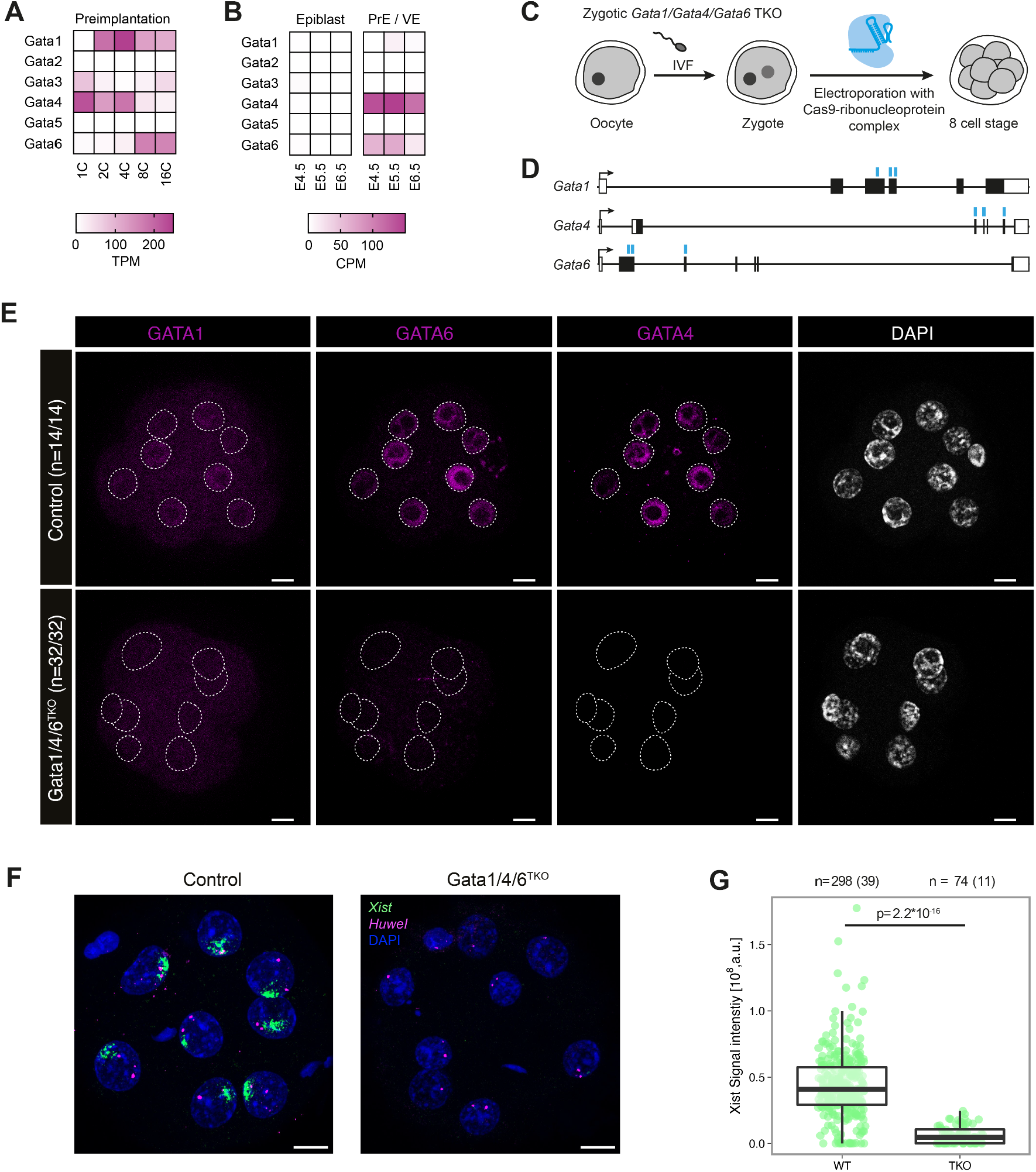
GATA factors are required for initial *Xist* upregulation *in vivo*. **(A-B)** Expression of GATA TFs during early development assessed by scRNA-seq (41,56). PrE primitive endoderm; VE visceral endoderm. **(C-G)** Zygotic triple knock-out (TKO) of *Gata1, Gata4* and *Gata6*. (C) Schematic depiction of the experimental workflow, where zygotes, generated by *in vitro* fertilization (IVF) were electroporated with Alt-R CRISPR/Cas9 ribonucleoprotein complex pre-assembled with 3 crRNAs targeting the *Gata1, Gata4* and *Gata6* coding sequences. Embryos were allowed to develop to the 8C stage. **(D)** Schematic depiction of *Gata1, Gata4* and *Gata6* genomic loci with regions targeted by crRNAs shown as blue lines. **(E)** Staining of the indicated GATA TFs. Dashed lines represent the nuclei as detected by DAPI staining. For the numbers indicated, two biological replicates were merged. **(F-G)** RNA-FISH for *Xist* and the X-linked *Huwe1* gene (nascent transcript) at the 8-cell stage. Only female embryos (2 *Huwe1* signals) were included in the analysis. In (G) the summed fluorescence intensity within the automatically detected *Xist* cloud is shown for individual cells. Embryos from two biological replicates were pooled. Statistical comparison was performed with a Wilcoxon ranksum test. The central mark indicates the median, and the bottom and top edges of the box indicate the first and third quartiles, respectively. The top and bottom whiskers extend the boxes to a maximum of 1.5 times the interquartile range; cell (embryo) numbers are indicated on top. The scale bars in (E+F) represent 10 μm

To investigate whether GATA factors might indeed be involved in *Xist* regulation during early embryogenesis, we tested whether the GATA-responsive enhancer element RE79 is part of the *tg80* and *tg53* single-copy transgenes, which can drive *Xist* expression in preimplantation embryos, but not in somatic cells (17,18). RE79 is located around the telomeric end of the transgenes, but the precise extent has never been mapped (Suppl. Fig. 7A). We therefore performed quantitative PCR on genomic DNA from mESCs derived from the *tg80* and *tg53* mouse lines. We found that RE79 is indeed part of *tg80* and *tg53* (Suppl. Fig. 7B), which might thus allow GATA factors to drive *Xist* expression from the transgene.

Next, we tested whether RE79 was required for maintenance of imprinted *Xist* expression. To this end, we introduced a ~500 bp deletion of RE79 in XEN and TS cells by Cas9-mediated gene editing (Suppl. Fig. 8). However, no consistent effect on *Xist* expression could be observed. The CUT&Tag data we had generated for H3K27ac in XEN and TS cells (Fig. 4A) suggested that the region upstream of *Xist* contains a number of additional active enhancer elements. These might act redundantly with RE79, which could explain why RE79 deletion did not affect *Xist* expression in these cell lines.

If robust maintenance of imprinted *Xist* expression is ensured through redundant regulation once iXCI is established, we reasoned that it might be easier to observe GATA-dependent *Xist* regulation at the onset of XCI. We therefore aimed to disrupt this putative mode of regulation during preimplantation development through zygotic electroporation of a Cas9 ribonucleoprotein complex. We initially tried to induce a RE79 deletion, but were unable to establish a robust genotyping strategy due to the limited material available. We therefore decided to delete selected GATA factors instead, where genotyping can be performed by protein staining.

We generated triple knock-out embryos of Gata1, Gata4 and Gata6 (Gata1/4/6^TKO^), as these factors exhibited high expression levels during the first days of development (Fig. 5A-D). When assaying for GATA1/4/6 protein expression at the 8-cell stage, we found that the KO strategy was highly efficient. All 32 Gata1/4/6^TKO^ embryos analyzed were deficient for all 3 factors, which were robustly detected in embryos electroporated with a control sgRNA targeting GFP (Fig. 5E). We therefore assayed *Xist* expression by RNA-FISH also at the 8-cell stage, where normally prominent *Xist* “clouds” covering the X chromosome are detected. Due to a developmental delay in the mutant embryos, less Gata1/4/6^TKO^ embryos could be analyzed than controls. We nevertheless observed a striking phenotype in the Gata1/4/6^TKO^ embryos, which showed generally very weak *Xist* signals and even absence of *Xist* upregulation in a subset of cells (Fig. 5F, Suppl. Fig. 9A). Quantification of *Xist* signals through automated image analysis revealed that *Xist* signal intensity was strongly reduced compared to control embryos (Fig. 5G). GATA factors are thus required for the initial upregulation of *Xist* after fertilization. Given the strong reduction of *Xist* expression upon loss of GATA TFs, the absence of GATA factors in the pluripotent epiblast (Fig. 5B) might contribute to *Xist* downregulation at that stage. With the GATA family we have therefore identified the first essential tissue-specific Xist activators and propose a key role for them in governing the initiation of XCI *in vivo*.

## Discussion

In this work, we identify GATA TFs as potent Xist activators and reveal a central role of GATA-mediated *Xist* regulation during early development. We show that all six family members are able to induce ectopic *Xist* upregulation in mESCs. We identify a distal enhancer element that mediates GATA6-dependent *Xist* induction and is bound by different GATA factors in extraembryonic lineages. Finally, we demonstrate that *Xist* upregulation is strongly impaired upon simultaneous deletion of three GATA TFs in mouse zygotes. Given that different subsets of GATA TFs are present in all Xist-expressing cells in preimplantation embryos, but absent from pluripotent cells, where *Xist* is downregulated, we propose a role for this TF family in controlling XCI patterns during early development.

From our results a new picture emerges of how XCI is regulated during early development. It has previously been suggested that the XCI pattern is mostly controlled through *Xist* repression by pluripotency factors, either through direct binding of a regulatory element within *Xist’s* intron 1, or indirectly through activation of *Xist’s* repressive antisense transcript *Tsix* (19,20,31,58). However, *Tsix* is not required for *Xist* repression in the epiblast (22,59) and deletion of the intron 1 binding site alone or in combination with a *Tsix* mutation does not lead to de-repression of *Xist* in mESCs (60,61). In light of our findings, these results can be explained by the absence of activating factors in mESCs. We demonstrate that GATA factors are needed for the first upregulation of *Xist* upon fertilization. Due to the fact that GATA TFs are expressed in a variety of combinations during preimplantation development and in extraembryonic lineages, they almost certainly contribute to the maintenance of *Xist* expression in those cellular contexts. The only cell type in the preimplantation embryo that does not express any GATA TF are pluripotent epiblast cells (62–64). Here, loss of GATA expression coincides with *Xist* downregulation and X-chromosome reactivation (8). Our finding that at least GATA1 is a strong *Xist* activator, when over-expressed in pluripotent stem cells, suggests that the loss of GATA expression is likely required to *Xist* downregulation. Since GATA factors are expressed in a wide variety of cell types, including the blood and the heart (43), this mode of regulation might also be involved in maintaining *Xist* expression in somatic cells.

We have identified a single enhancer element, namely RE79, located ~100 kb upstream of the *Xist* promoter, that mediates GATA-induced *Xist* upregulation. We have recently shown that this element does not control *Xist* at the onset of rXCI (16). A different set of long-range elements governs *Xist* upregulation in the context of rXCI, which are bound by TFs associated with the pluripotent state in postimplantation embryos, such as OTX2 and SMAD2/3 (16). Tissue-specific expression of *Xist* thus appears to be orchestrated by a series of distal enhancer elements, which respond to lineage-specific transcription factors, such as GATA4 and GATA6 in the primitive endoderm, GATA2 and GATA3 in the trophectoderm and OTX2 and SMAD2/3 in the epiblast. These long-range elements can, however, only induce *Xist* expression, if the promoter-proximal region is not repressed either by the rodent-specific imprint or through the RNF12-REX1-axis, which helps prevent *Xist* upregulation in male cells.

Imprinted XCI in extraembryonic tissues has evolved specifically in rodents. However, also in human embryos *Xist* is upregulated shortly after fertilization (65). In contrast to mice, *Xist* is expressed from all X chromosomes in male and female preimplantation embryos, but does not yet initiate XCI (66,67). Given that multiple GATA transcription factors are expressed during preimplantation development in human embryos (67–69), it is tempting to speculate that biallelic *Xist* upregulation is a result of GATA-dependent activation that can act on both X chromosomes, as the maternal *Xist* locus is not imprinted in humans.

A commonly assumed regulatory principle is that ubiquitous expression is governed by broadly expressed TFs (70). Our results unveil a conceptually different regulatory strategy for ubiquitous expression: Members of a TF family are expressed in specific cell types, yet together covering many different tissues. In this way, a group of TFs with tissue-specific expression patterns, but overlapping DNA binding preferences, would jointly drive near-ubiquitous expression of a target gene. Ongoing efforts to precisely map the transcriptome across tissues, such as the human cell atlas, will allow us to understand how common this regulatory strategy is used to shape gene expression in complex organisms.

## Methods

### Cell lines

The female TX1072 (clone A3), TX-SP106 (Clone D5) and TX-SP107 (Clone B6) mESC lines as well as the male E14-STN_ΔTsixP_ mESC cell line were described in (16). Briefly, the female TX1072 cell line (clone A3) is a F1 hybrid ESC line derived from a cross between the 57BL/6 (B6) and CAST/EiJ (Cast) mouse strains that carries a doxycycline-responsive promoter in front of the *Xist* gene on the B6 chromosome. TX1072 XO (clone H7/A3) is an XO line that was subcloned from TX1072 and has the B6 X chromosome. The TX-SP106 (Clone D5) mESC line stably expresses PYL1-VPR-IRES-Blast and ABI-tagBFP-SpdCas9, constituting a two-component CRISPRa system, where dCas9 and the VPR activating domain are fused to ABI and PYL1 proteins, respectively, which dimerize upon treatment with abscisic acid (ABA). The TX-SP107 (Clone B6) mESC line stably expresses PYL1-KRAB-IRES-Blast and ABI-tagBFP-SpdCas9, constituting a two-component CRISPRi system, where dCas9 and the KRAB repressor domain are fused to ABI and PYL1 proteins, respectively, which dimerize upon ABA treatment. Since repression in TX-SP107 cells transduced with sgRNAs was often observed already without ABA treatment, we could not make use of the inducibility of the system. Instead, TX-SP107 cells were always treated with ABA (100 μM) 72 h before the analysis and effects were compared to NTC sgRNAs. The male E14-STN_ΔTsixP_ mESC cell line expresses the CRISPRa SunTag system (27,28) under a doxycycline-inducible promoter and carries a 4.2 kb deletion around the major *Tsix* promoter (ChrX:103445995-103450163, mm10).

Female XEN cells were a kind gift from the Gribnau lab (71). XEN XX #12 cell line was derived from a crossing of C57BL/6 (B6) female mice with CAST/Eij (Cast) males. NGS karyotyping detected trisomies of chr. 1, 14 and 16. A CD1-derived female TSC line was a kind gift from the Zernicka-Goetz lab. Low-passage Hek293T cells were a kind gift from the Yaspo lab. Details on all cell lines are given in Suppl. Table 4. All cell lines were routinely checked for XX status via RNA-FISH using a BAC probe for *HuweI* as described below.

### mESC culture and differentiation

All mESC lines were grown on 0.1% gelatin-coated flasks in serum-containing medium (DMEM (Sigma)), 15% ESC-grade FBS (Gibco), 0.1 mM β-mercaptoethanol), either supplemented with 1000 U/ml leukemia inhibitory factor (LIF, Millipore) only (E14-STN_ΔTsixP_, TX-SP106) or with LIF and 2i (3 μM Gsk3 inhibitor CT-99021, 1 μM MEK inhibitor PD0325901, Axon) (TX-SP107, TX-SP107-ERT2-Gata6-HA). Differentiation was induced by LIF or LIF/2i withdrawal in DMEM supplemented with 10% FBS and 0.1 mM β-mercaptoethanol at a density of 4-4.2*10^4^ cells/cm^2^ in fibronectin-coated (10 μg/ml) tissue culture plates.

In CRISPRa-SunTag (E14-STN_ΔTsixP_) experiments, the cells were treated with doxycycline (1 μg/ml) for 3 days before harvesting. In CRISPRi and CRISPRa-VPR (TX-SP106) experiments, the cells were treated with Abscisic acid (ABA, Sigma 100 μM) for 3 days before harvesting. For nuclear translocation of ERT2-Gata6-HA, the cells were treated with 4-Hydroxytamoxifen (4OHT, Sigma, 2.5 μM).

### XEN and TS cells culture

Female XEN cell line was grown on 0.2% gelatin-coated flasks following the Rossant lab XEN stem cell protocol (https://lab.research.sickkids.ca/rossant/wp-content/uploads/sites/12/2015/08/XEN-Stem-Cell-protocol1.pdf) in serum-containing XEN medium (RPMI 1640 (Sigma, M3817)), 15% ESC-grade FBS (Gibco), 0.1 mM β-mercaptoethanol (Sigma), 1 mM sodium pyruvate (Gibco) and 2 mM L-glutamine (Life technologies).

Female TSCs were grown on MEFs in serum containing TSC medium (RPMI, 20 % fetal bovine serum, 1 mM sodium pyruvate, 100 mM 2-mercaptoethanol, 50 μg/ml penicillin/streptomycin and 2 mM L-Glutamine; FGF4 (25 ng/ml, R&D System) and Heparin (1 μg/ml, Sigma) were added to the medium fresh prior to each use) (10). Before sample collection, TSCs were passaged at least twice without MEFs to dilute out feeder cells. During this time cells were cultured in MEF-conditioned medium (70% MEF-conditioned medium, 30% TSC medium, FGF4 (37.5 ng/ml, R&D System), Heparin (1.5 μg/ml, Sigma)).

### Generation of transgenic cell lines

Transgenic cell lines were generated via lentiviral transduction. To package lentiviral vectors into lentiviral particles, 1*10^6^ HEK293T cells were seeded into one well of a 6-well plate and transfected the next day with the lentiviral packaging vectors: 1.2 μg pLP1, 0.6 μg pLP2 and 0.4 μg pVSVG (Thermo Fisher Scientific), together with 2 μg of the desired construct using Lipofectamine 2000 (Thermo Fisher Scientific). HEK293T supernatant containing the viral particles was harvested after 48 h. 0.1-0.2*10^6^ mESCs were seeded per well in a 24-12-well plate in conventional ESC medium and transduced the next day with 0.25-0.5 ml of 10:1 concentrated (lenti-X, Clontech) supernatant with 8 ng/μl polybrene (Sigma Aldrich). Transgenic cells were selected with puromycin (sgRNA plasmids) (1 ng/μl, Sigma) or hygromycin (FIREWach plasmids, 200 ng/μl, VWR) starting two days after transduction. Selection was kept for the entire experiment.

Cell lines overexpressing Gata1-6, Xist, Esx1, Cdx4 and Nup62cl via the CRISPRa SunTag system were generated by lentiviral transduction of E14-STN_ΔTsixP_ cells with sgRNAs, as indicated in the respective figure legend, targeted to the respective promoters or non-targeting controls (Suppl. Table 4).

TX-SP107 CRISPRi cell lines for Gata1, Xist and Gata REs (RE57/RE79/RE97) were generated by lentiviral transduction of TX-SP107 (Fig. 2E-G)/TX-SP107-ERT2-Gata6-HA (Fig. 5G+H) cells, carrying an ABA-inducible dCas9-KRAB system with plasmids carrying 1 (Xist) or 3-4 (Gata1/REs) sgRNAs targeted to the respective genomic loci or non-targeting controls, (SP125_LR249, SP199_mgLR9, SP199_mgLR22/23, SP199_mgVS012, SP199_mgLR15/16/17).

Cell lines expressing the FIREWach reporter plasmid (55) with the Gata REs regions and over-expressing Gata6 via the CRISPRa-ABA-inducible VPR system were generated by two rounds of lentiviral transduction. First, TX-SP106 (Clone D5) cells were transduced with plasmids carrying multi-sgRNAs targeting the *Gata6* promoter or non-targeting controls, (SP199_mgLR7, SP199_mgLR15/16). Then, either the empty (SP307) or the RE-containing FIREWach plasmids (SP379, SP376, SP418) were lentivirally integrated into the cells, which were treated with abscisic acid (ABA, Sigma 100 μM) for 3 days before harvesting.

### Genome Engineering

#### Generation of RE79 knock-out XEN cell lines

To analyze the role of RE79 in in maintaining Xist expression in XEN cells, a heterozygous RE79 deletion was introduced at the inactive X-chromosome (Cast) in a hybrid female XEN cell line (B6xCast) and the effect on Xist expression was analyzed. To this end, 2 guide RNAs (crRE79_1/crRE79_2) were used with the Alt-R® CRISPR-Cas9 System (IDT), which contains all necessary reagents for the delivery of Cas9-gRNA ribonucleoprotein complexes (RNP) into target cells. Briefly, crRNAs and tracrRNA-atto550 were mixed in equimolar concentrations and the 2 crRNAs and tracrRNA duplexes were subsequently pooled together. 2.1 μl PBS, 1.2 μl of the tracr duplex (100 μM Stock), 1.7 μl Cas9 (61 μM Stock) and 1 μl electroporation enhancer were pipetted together and incubated for 20 min. 10^5^ cells were nucleofected with the mixture using the DA113 program of the Amaxa 4D-Nucleofector (Lonza) and plated on a 0.2% gelatin-coated 24-well plate. After 5 days, cells were seeded in a 96-well plate on a feeder layer at a low density (~ 1 cell per well) in XEN media supplemented with 10 μM ROCK inhibitor Y-27632 (Tocris #1254) until clones appeared (approximately 5 days). Individual clones were expanded and genotyped for the presence of the RE79 deletion as outlined below.

#### RE79 knock-out in TSCs

To analyze the role of RE79 in maintaining Xist expression in TS cells, a homozygous RE79 deletion was introduced in a female TS line, using the sgRNA/Cas9 system. TSCs were transfected with two PX458 plasmids (Addgene, #48138) each containing one single guide RNA (RE79_1/RE79_2) that together allow to cut out the locus of interest. TSCs were transfected using FuGENE HD Transfection Reagent (Promega). Briefly, 3*10^5^ cells were plated the day before transfection (feeder free) in MEF-conditioned medium (supplemented with 37.5 ng/ml FGF4 and 1.5 μg/ml Heparin). On the day of transfection 8 μg of plasmid DNA was diluted in 125 μl Opti-MEM (Thermofisher) and 25 μl of FuGENE Reagent was diluted with 100 μl Opti-MEM. Diluted FuGENE was added to diluted DNA, incubated at room temperature for 15 min and added to the cells dropwise. Medium was changed on the next day and GFP positive cells were sorted 48-72 h post transfection and plated on feeder cells in standard TSC medium containing 10 μM ROCK inhibitor Y-27632 (Tocris #1254). Clonal TSC colonies were picked 7-10 days post sort, expanded without feeder cells in MEF-conditioned medium with ROCK inhibitor. Once confluent, the clones were split into two plates for expansion and genotyping, respectively, and were expanded without feeder cells in MEF-conditioned medium with ROCK inhibitor.

#### Genotyping of engineered XEN and TS clones

For genotyping both XEN and TSC RE79 KO clones, gDNA was isolated from a 96-well plate. The cells were washed with PBS and lysed in the 96-wells plate with 50 μl Bradley lysis buffer (10 mM Tris-HCl pH 7.5, 10 mM EDTA, 0.5% SDS, 10 mM NaCl, 1 mg/ml Proteinase K (Invitrogen)). The plate was incubated overnight at 37-55°C in a humidified chamber. To precipitate gDNA, 150 μl ice-cold 75 mM NaCl in 99% EtOH was added per well and the plate was incubated for 0.5-4 h at room temperature. The plate was centrifuged for 15 min at 4000 rpm and 4 °C. The pellet was washed 1-3 times with 70% EtOH and centrifuged for 15 min at 4000 rpm and air dried for 10 min at 45°C. The gDNA was resuspended in 200 μl water (XEN) or 30 μl EB (TSC) for 2 h at 45 °C. The clones were characterized by PCR with the primers ASK259/ASK260 that can detect both WT and deleted alleles. A small number of positive clones were expanded and PCR genotyping was repeated on gDNA isolated using the DNeasy Blood and Tissue Kit (Qiagen) or GeneJET Genomic DNA Purification Kit (Thermo). To identify the targeted allele in XEN heterozygous KO clones, amplicons containing SNPs (PCR amplified with primers ASK259/LR588) were gel-purified and sequenced with the primer LR588.

#### Generation of Gata1/Gata4/Gata6 triple knock-out mouse embryos

All animal procedures were conducted as approved by the local authorities (LAGeSo Berlin) under the license number G0243/18-SGr1. Oocytes were obtained from donor B6D2F1 female mice of 7-9 weeks of age (Envigo) by superovulation; hormone priming with 5 IU of PMSG followed by 5 IU of HCG 46 h later. 12 h after hormone priming, MII stage oocytes were isolated and cultured in standard KSOM media. Zygotes for knockout experiments were obtained by performing *in vitro* fertilization (IVF) with donor oocytes and sperm under standard conditions. Sperm used for IVF is prepared from fertile F1 males (B6/CAST) as previously described (72). Electroporation was performed as previously described (72) with pre-assembled Alt-R CRISPR/Cas9 Ribonucleoprotein complex (IDT). Three guides targeting exons were used for every target gene. Guide RNA sequences used can be found in Suppl. Table 4. Zygotes electroporated with a mock guide (targeting GFP) were used as control. Electroporated embryos were washed and cultured in KSOM medium *in vitro* under standard conditions (5% CO_2_, 37 °C). Gata1/4/6^TKO^ embryos developed more slowly compared to the controls.

### Flow cytometry

For flow RNA fluorescence in-situ hybridization (Flow-FISH) the PrimeFlow RNA assay (Thermofisher) was used according to the manufacturer’s recommendations. Specifically, the assay was performed in conical 96-well plates with 5*10^6^ cells per well with *Xist*-specific probes, labeled with Alexa-Fluor647 (VB1-14258) (Thermo Fisher Scientific). Samples were resuspended in PrimeFlow™ RNA Storage Buffer before flow cytometry.

Flow cytometry data was collected using a BD FACSAria II, BD FACSAria Fusion or BD FACS Celesta flow cytometer. The sideward and forward scatter areas were used to discriminate cells from cells’ debris, whereas the height and width of the sideward and forward scatter were used for doublet discrimination. At least 30,000 cells were measured per sample. FCS files were analyzed using RStudio with the *flowCore* (v1.52.1) and *openCyto* packages (v1.24.0) (73,74).

For Flow-FISH, all cells that showed a fluorescence intensity above the 99th-percentile of the undifferentiated cell population control, which does not express *Xist*, were marked as *Xist*-positive. These cells were then used to calculate the geometric mean in the *Xist*-positive fraction after background correction by subtracting the geometric mean of the undifferentiated control. In the enhancer-reporter assay, the geometric mean of the GFP fluorescence intensity was calculated and background-corrected by subtracting the geometric mean of the TX-SP106 non-transduced control (GFP negative).

### Molecular Cloning

#### sgRNA cloning

To facilitate diagnostic digestion after cloning, an AscI restriction site was added to the original pU6-sgRNA-EF1*a*-puro-T2A-BFP plasmid (Addgene #60955, (75)) between the BlpI and BstXI sites, resulting in plasmid SP125, by annealing the oligos LR148/LR149 that contain the restriction site. Single sgRNAs for CRISPRa were cloned into a BlpI and BstXI digested pU6-sgRNA-EF1*a*-puro-T2A-BFP plasmid by annealing oligos containing the guide sequence and recognition sites for BlpI and BstXI (Oligo F: 5’TTGGNNN…NNNGTTTAAGAGC3’and Oligo R: 5 ’TTAGCTCTTAAAC NNN…NN NCCAACAAG 3’) and ligating them together with the linearized vector using the T4 DNA ligase enzyme (NEB). Cloning of sgRNAs in a multiguide expression system (SP199) was performed as described previously (38). Briefly, three or four different sgRNAs targeting the same gene/RE (Suppl. Table 4) were cloned into a single sgRNA expression plasmid with Golden Gate cloning, such that each sgRNA was controlled by a different Pol III promoter (mU6, hU6 hH1, h7SK) and fused to the optimized sgRNA constant region (76). The vector (SP199) was digested with BsmBI (New England Biolabs) 1.5 h at 55 °C and gel-purified. Three fragments containing the optimized sgRNA constant region coupled to the mU6, hH1 or h7SK promoter sequences were synthesized as gene blocks (IDT). These fragments were then amplified with primers that contained part of the sgRNA sequence and a BsmBI restriction site (primer sequences can be found in Suppl. Table 4) and PCR-purified using the gel and PCR purification kit (Macherey & Nagel). The vector (100 ng) and two (for cloning 3 sgRNAs) or three (for cloning 4 sgRNAs) fragments were ligated in an equimolar ratio in a Golden Gate reaction with T4 ligase (New England Biolabs) and the BsmBI isoschizomer Esp3I (New England Biolabs) for 20 cycles (5 min 37°C, 5 min 20°C) with a final denaturation step at 65 °C for 20 min. Vectors were transformed into NEB Stable competent E.coli. Successful assembly was verified by ApaI digest and Sanger sequencing.

#### ERT2-Gata6-HA-T2A-Hygro overexpression construct

The plasmid was generated by standard molecular cloning techniques and its sequence is provided in the supplemental material (Suppl. Table 5). In brief, to generate ERT2-Gata6-HA-T2A-Hygro (SP299), the backbone of pLenti-ERT2-FLAG-Gal4-NLS-VP16-P2A-Pur o (SP265) was used. SP265 was digested with NdeI/MluI (New England Biolabs) to remove FLAG-Gal4-NLS-VP16-P2A-Puro. The backbone was ligated with a HA-T2A-HygroR fragment, that was amplified from lenti-MS2-p65-HSF1_Hygro plasmid (Addgene #61426) using a primer that contained the HA-tag sequence via InFusion cloning. *Gata6* cDNA was PCR amplified from pSAM2_mCherry_Gata6 (Addgene #72694) and ligated using InFusion cloning.

#### Cloning of the Gata REs into the FIREWACh enhancer plasmid

To generate RE-containing enhancer reporter plasmids, each RE (RE57, RE79 and RE97) was PCR-amplified from BAC (RP23-11P22, RP23-423B1) or genomic DNA with overhangs for InFusion cloning (Takara). The fragments were ligated into a BamHI digested FIREWACh plasmid FpG5 (Addgene #69443) (55), to yield plasmids SP379, SP376, SP418.

FIREWACh RE In-Fusion cloning (Takara) was carried out in a 2:1 insert/vector ratio. Plasmid sequences are given in Suppl. Table 5.

### RNA extraction, reverse transcription, qPCR

For gene expression profiling, cells were washed and lysed directly in the plate by adding 500 μl of Trizol (Invitrogen). RNA was isolated using the Direct-Zol RNA Miniprep Kit (Zymo Research) following the manufacturer’s instructions with on-column DNAse digestion. For quantitative RT-PCR (qRT-PCR), up to 1 μg RNA was reverse transcribed using Superscript III Reverse Transcriptase (Invitrogen) with random hexamer primers (Thermo Fisher Scientific) and expression levels were quantified in the QuantStudio™ 7 Flex Real-Time PCR machine (Thermo Fisher Scientific) using Power SYBR™ Green PCR Master Mix (Thermo Fisher Scientific) normalizing to Rrm2 and Arp0. Primers used are listed in Suppl. Table 4.

#### RNA FISH on embryos

To prepare preimplantation embryos (8 C stage) for RNA-FISH, embryos were washed through a series of KSOM drops (Sigma), followed by a series of Tyrode’s solution. Zona pellucida was removed by incubating the embryos in Tyrode’s solution (Sigma) for 10-30 sec until the zona was dissolved. The embryos were washed through a series of PBS + 0.4% BSA prior to mounting onto Poly-L-Lysine (Sigma) coated (0.01% in H_2_O, 10 min incubation at room temperature) coverslip #1.5 (1 mm). Embryos were allowed to attach for about 2 min after which excess volume was removed and allowed to dry for 30 min. Embryos were fixed in 3% paraformaldehyde in PBS for 10 min at room temperature and permeabilized for 4 min on ice in PBS containing 0.5% Triton X-100 and 2 mM Vanadyl-ribonucleoside complex (New England Biolabs). Coverslips were stored in 70% EtOH in −20°C no longer than 1 day before further processing.

RNA-FISH was performed using the plasmid probe *p510* spanning the genomic sequence of *Xist* and the BAC probe (RP24-157H12) for *Huwe1* as described previously with minor modifications (77). Both probes were labeled by nicktranslation (Abbot) with dUTP-Green (Enzo) or dUTP-Atto550 (Jena Bioscience), respectively. Per coverslip, 120-200 ng of each probe were ethanol precipitated (Cot1 repeats were included for *Huwe1* in order to suppress repetitive sequences in the BAC DNA that could hamper the visualization of specific signals), resuspended in 3-6 μl formamide and denatured (10 min 75 °C). For *Huwe1*, a competition step of 1 h at 37 °C was added. Before incubation with the probe, the samples were dehydrated through an ethanol series, 90% and 100%, twice of each (5 min each wash), and subsequently air-dried. Probes were hybridized in a 12 μl hybridization buffer overnight (50% Formamide, 20% Dextran Sulfate, 2x SSC, 1 μg/μl BSA, 10 mM Vanadyl-ribonucleoside). To reduce background, three 5 min washes were carried out in 50% Formamide/2x SSC (pH 7.2) and one 5 min wash in 2x SSC at 42 °C. Two additional washes in 2x SSC were carried out at room temperature and 0.2 mg/ml DAPI was added to the first wash. The samples were mounted using Vectashield mounting medium (Vector Laboratories).

Embryo image acquisition was performed using an inverted laser scanning confocal microscope with Airyscan (LSM880, Zeiss) with a 63x/1.4 NA oil objective, lateral resolution of 0.07 μm and 0.28 μm Z-sections in Fast Airyscan mode. Acquisition was performed under Zeiss ZEN 2.6 black software.

#### Automated analysis of RNA-FISH in embryos

Confocal Z-Stacks were 3D airyprocessed using ZEN 2.6 Black and all subsequent analyses were performed in ZEN 3.2 or Zen 3.4 blue (both Zeiss, Germany) equipped with the Image Analysis module. The sex of each embryo was determined visually based on the RNA-FISH signal for the nascent transcript for *Huwe1*, an X-linked gene that is not yet silenced by XCI at the stages analyzed (2 signals per nucleus in females, one in males). Only female embryos were included in the analysis. Images were maximum intensity projected and a spot detector was used to identify primary objects (nuclei) by Gaussian smooth, Otsu-thresholding, dilation and water shedding. The resulting objects were filtered by area of 100-450 μm^2^ and circularity (sqrt((4*area)/(π*FeretMax^2^))) of 0.7-1. *Xist* clouds were identified as a subclass within primary objects. Here, images were smoothed, background-subtracted (rolling ball), followed by a fixed intensity threshold to identify spots. Only nuclei with a Huwe1 signal were included in the downstream analysis. The summed signal intensity within the identified Xist spots were compared between cells in wildtype and TKO embryos using a Wilcoxon ranksum test. Since the TKO embryos exhibited a developmental delay, less 8-cell embryos could be analyzed compared to the control.

### Immunofluorescence combined with RNA FISH (IF-RNA-FISH)

IF-RNA-FISH was performed according to the Stellaris protocol for adherent cells, https://www.protocols.io/view/Stellaris-RNA-FISH-Sequential-IF-FISH-in-Adherent-ekzbcx6 with minor modifications. TX-SP107-ERT2-Gata6-HA cells were grown under 2i/LIF conditions. Two days before fixation, the cells were plated on fibronectin-coated coverslips (18 mm, Marienfeld) at a density of 2*10^4^ cells cm^−2^ in medium without 2i, which helps cells to spread sufficiently for imaging. Cells were fixed in 3% paraformaldehyde in PBS for 10 min at room temperature and permeabilized for 5 min at room temperature in PBS containing 0.1% Triton X-100, after 6 h and 24 h of 2.5 μM 4OHT treatment as applicable. The coverslips were incubated with an HA-specific antibody (Abcam, ab9110 1:1000) in PBS for 1 h at room temperature, then washed three times for 10 min with PBS, followed by a 1 h incubation with an Alexa-555 labeled Goat anti-rabbit antibody (Invitrogen A-21428, 0.8 μg ml^−1^). After three washes, the cells were fixed again with 3% paraformaldehyde in PBS for 10 min at room temperature, followed by three short washes with PBS and two washes with 2x SSC. *Xist* was detected using Stellaris FISH probes (Biosearch Technologies). Coverslips were incubated for 5 min in wash buffer containing 2x SSC and 10% formamide, followed by overnight hybridization at 37 °C with 250 nM of FISH probe in 50 μl Stellaris RNA FISH Hybridization Buffer (Biosearch Technologies) containing 10% formamide. Coverslips were washed twice for 30 min at 37 °C with 2x SSC/10% formamide with 0.2 mg/ml Dapi being added to the second wash. Prior to mounting with Vectashield mounting medium coverslips were washed with 2x SSC at room temperature for 5 min. Details on the antibodies and probes used are found in Suppl. Table 4.

Cell images were acquired using a widefield Axio Observer Z1/7 microscope (Zeiss) using a 100x Oil Immersion objective (NA=1.4). Image analysis was carried out using Zen 3.1 blue (Zeiss). For each sample and replicate 5 tile regions were defined, the optimal focus was adjusted manually. The focused image was used as a center for a z-stack of 62 slices with an optimal distance of 0.23 μm between individual slices. Thereby, a total stack height of 14.03 μm was achieved covering slightly more than the cell height to ensure capturing of all events.

#### Automated analysis of IF-RNA-FISH

Image analysis was performed with ZEN 3.2 and 3.4 (Carl Zeiss, Germany). Images underwent a Maximum Intensity Projection (MIP) of the full Z-Stack of 62 slices. Segmentation of DAPI stained nuclei was achieved with a priori trained Intellesis model. The identified objects were only kept in the subsequent steps, if they exhibited a circularity (Sqrt(4 × Area / π × FeretMax^2^)) of 0.5-1 and an area of 50-300 μm^2^. Around each nucleus a ring (width 30 pix = 2.64 μm) was drawn and used as a surrogate for the cytoplasmic region. From the nuclear and cytoplasmic compartments the mean fluorescence intensity was extracted for the Gata6-HA staining and the nuclear-to-cytoplasmic ratio was calculated as a proxy for nuclear translocation. For identification of nuclear *Xist* signals, images were Gaussian smoothed, followed by a rolling ball background subtraction (radius 20 pixel) and a fixed intensity threshold. The identified areas were filtered to fit a circularity between 0.5-1. All cells with >2 *Xist* objects were excluded from the analysis.

### Immunofluorescence staining

Embryos were washed through a series of KSOM drops (Sigma), followed by a series of PBS + 0.4% BSA. Fixation was performed by incubation with 4% PFA for 15 mins. PFA was washed off by a series of washes in PBS + 0.5% TritonX-100 (PBS-T). Embryos were permeabilized in PBS-T for 20 min at room-temperature. After permeabilization, samples were washed in PBS-T and blocked in PBS-T + 2% BSA + 5% goat serum for 1 h at room temperature. Primary antibodies were diluted in blocking buffer (PBS-T + 2% BSA + 5% goat serum) overnight at 4°C. Following incubation with the primary antibody, samples were washed three times for 10 min at room temperature in PBS-T + 2% BSA and subsequently incubated with secondary antibodies (1:1000) in PBS-T + 2% BSA + 5% goat serum for 1 h at room temperature. Samples were washed three times 10 min at room temperature in PBS-T. After the last washing step, embryos were transferred to mounting medium (Vectashield, H1200) and further to a glass slide (Roth) and sealed with a cover glass (Brand, 470820). Detailed information on the antibodies used is given in Suppl. Table 4. Images were acquired with ZEISS LSM880 microscope at 40x magnification. Images were processed with ImageJ. Background fluorescence was subtracted by using *rolling ball radius* method (ImageJ) with 50 pixels as threshold.

### PyroSequencing

To quantify relative allelic *Xist* expression for XEN RE79 clones, an amplicon containing a SNP at the Cast allele was amplified by PCR from cDNA using Hot Start Taq (Qiagen) for 38 cycles. The PCR product was sequenced using the Pyromark Q24 system (Qiagen). Assay details are given in Suppl. Table 4.

### Tg80 mapping

QPCR was performed on genomic DNA from IKE15-9TG80 and IKE14-2TG53 (XY-tg), carrying a single copy of YAC PA-2 (18) and E14-STN_ΔTsixP_ (reference XY DNA) using primer pairs detecting different positions within the *Ftx* genomic locus. QPCR measurements were normalized to amplification from an X-linked locus outside of the YAC region (LR621/622). By calculating the ratio of the relative expression between the two cell lines, each genomic position could be classified as either internal (ratio ~2) or external (ratio ~1) to the YAC region.

### CRISPRa screen

#### CRISPRaX sgRNA library design

To target protein- and non protein-coding X-linked genes via CRISPRa, sgRNA sequences were extracted from the mouse genome-wide CRISPRa-v2 library (78) and complemented with newly-designed sgRNAs using the CRISPR library designer (CLD) software (79). Using Ensembl release (corresponding to genome assembly mouse mm10) and FANTOM 5 CAGE data (80), a list of all TSSs for expressed genes (read count > 0, based on bulk-RNA seq data for female mESCs in 2i/LIF and 36 h −2i/LIF conditions) was compiled. All newly-designed sgRNAs were *in-silico* tested for off-target effects in other promoter regions (550 bp window upstream of a TSS). In total the libraries targets 2695 TSSs on the X chromosome, corresponding to 757 genes. Each TSS was targeted by 6 sgRNAs in a window between 550 and 25 bp upstream of the TSS. In cases where two TSSs were in close proximity, the same guides were used to target different TSSs. Additionally, two verified sgRNAs for *Xist* and guides targeting a series of known X-linked *Xist* regulators (*Rnf12, Ftx, Jpx*), autosomal *Xist* regulators (*Nanog, Zfp42, Sox2, Myc, Klf4, Esrrb, Pou5f1, Prdm14, Ctcf, Yy1, Eed, Chd8, Kat8, Msl1, Msl2, Kansl3, Kansl1, Mcrs1, Dnmt1*) (19–23,31,32,46,81–85), and *Xist*-interacting proteins (*Spen, Lbr, Saf-A, Hnrnpk*) (86–88) were included in the CRISPRaX library as well as 200 non-targeting controls. The final library contained 8973 sgRNAs, which targeted 780 genes. The library composition is provided in Suppl. Table 1.

#### Cloning of CRISPRaX sgRNA library

The CRISPRaX sgRNA library was cloned into SP125, a modified pU6-sgRNA EF1Alpha-puro-T2A-BFP (pLG1) sgRNA expression plasmid (Addgene #60955, (75)) where an AscI restriction site was added between the BstXI and the BlpI sites that enabled diagnostic digestion after ligation for verification of positive colonies. The library was cloned following the Weissman lab protocol https://weissmanlab.ucsf.edu/CRISPR/Pooled_CRISPR_Library_Cloning.pdf. sgRNA sequences, G+19 nt, were synthesized by CustomArray flanked with OligoL (CTGTGTAATCTCCGACACCCACCTTGTTG) and OligoR (GTTTAAGAGCTAAGCTGGCCTTTGCATGTTGTGGA) sequences. For library amplification, 3 PCR reactions (Primer sequences in Suppl. table 4, LR169/LR170) with approx. 5 ng of the synthesized oligo pool were carried out using the Phusion High Fidelity DNA Polymerase (New England Biolabs), with a total of 15 cycles and an annealing temperature of 56 °C. The 3 PCR reactions were pooled and the 84 bp amplicons were PCR purified on a Qiagen Minelute column.

1 μg of the amplified sgRNAs was digested with BstXI (Thermo Fisher Scientific) and Bpu1102I (BlpI, Thermo Fisher Scientific) overnight at 37 °C. The digest was run on a 20% native acrylamide gel following staining with SYBR™ Safe DNA Gel Stain (Invitrogen) for 15 min. The 33 bp DNA fragment was extracted from the gel according to the Weissman lab protocol above. One 20 μl ligation reaction using T4 ligase (New England Biolabs) was carried out using 0.9 ng of the gel-purified insert and 500 ng of the vector. The reaction was EtOH-precipitated to remove excess salts which might impair bacterial transformation and resuspended in 20 μl H2O. 8 μl of the eluted DNA were transformed into 20 μl of electrocompetent cells (MegaX DH10B, Thermo Fisher Scientific) according to the manufacturer’s protocol using the ECM 399 electroporator (BTX). After a short incubation period (1 h, 37 °C 250 rpm) in 1 ml SOC medium, 9 ml of LB medium with Ampicillin (0.1 mg/ml, Sigma) were added to the mixture and dilutions were plated in Agar plates (1:100, 1:1000 and 1:10000) to determine the coverage of the sgRNA library (2000x). 500 ml of LB media with Ampicillin were inoculated with the rest of the mixture and incubated overnight for subsequent plasmid purification using the NucleoBond Xtra Maxi Plus kit (Macherey-Nagel) following the manufacturer’s instructions. To confirm library composition and even sgRNA representation by deep-sequencing a PCR reaction was carried out to add illumina adaptors and a barcode by using the Phusion High Fidelity DNA Polymerase (New England Biolabs), with an annealing temperature of 56 °C and 15 cycles (LR177/LR175, see Suppl. Table 4). The PCR amplicon was gel-purified by using the QIAquick Gel Extraction Kit (Qiagen) following the manufacturer’s instructions. Library was sequenced paired-end 75 bp on the HiSeq 4000 Platform using the sequencing primer LR176 yielding approximately 6 Mio. fragments. Read alignment statistics found in Suppl. Table 1).

#### Viral packaging of sgRNA library

To package the CRISPRaX library into lentiviral particles, HEK293T cells were seeded into 11 10 cm plates. The next day at 90% confluence each plate was transfected with 6.3 μg of pLP1, 3.1 μg of pLP2 and 2.1 μg of VSVG packaging vectors (Thermo Fisher Scientific) together with 10.5 μg of the CRISPRaX library plasmid in 1 ml of Opti-MEM (Life technologies) using 60 μl Lipofectamine 2000 reagent (Thermo Fisher Scientific) according to the manufacturer’s instructions. After 48 h the medium was collected and centrifuged at 1800 x g for 15 min at 4 °C. Viral supernatant was further concentrated 10-fold using the lenti-X^TM^ Concentrator (Takara Bio) following the manufacturer’s instructions and subsequently stored at −80 °C.

To assess the viral titer, 4 serial 10-fold dilutions of the viral stock were applied to each well of a 6-well mESC plate (MOCK plus 10^−3^ to 10^−6^) for transduction with 8 ng/μl polybrene (Merck). Two replicates were generated for each well. Selection with puromycin (1 ng/μl, Sigma) was started 2 days after transduction and colonies were counted after 7 days. The estimated titer was 5.43*10^6^ transducing units (TU) per ml.

#### Transduction

For the CRISPRa-SunTag screen, male E14-STN_ΔTsixP_ mESCs were passaged twice before 1.2*10^7^ cells were transduced with the CRISPRaX sgRNA library (MOI=0.3). Puromycin selection (1 ng/μl, Sigma) was started 48 h after transduction and kept until the end of the experiment. Four days after transduction, 7.2*10^7^ cells were differentiated by LIF withdrawal for 2 days. Expression of CRISPRa-SunTag system was induced using doxycycline (Clontech, 1 μg/ml) one day before differentiation and kept throughout the rest of the protocol. Cells were harvested with trypsin to reach a single cell suspension for Flow-FISH after 2 days of differentiation.

#### Flow-FISH and cell sorting

Phenotypic enrichment based on RNA levels was performed as previously described (89). The PrimeFlow RNA assay (Thermofisher) was used as described above. 2.4*10^8^ cells were stained, while 2*10^7^ cells were snap-frozen after the second fixation step to be used as the unsorted fraction. The 15% of cells with the highest fluorescence were sorted using a BD FACSAria II flow cytometer, recovering 7-15*10^6^ cells per replicate. After sorting, the cell pellet was snap-frozen and stored at −80 °C for further analysis.

#### Preparation of sequencing libraries and sequencing

Sequencing libraries were prepared from both sorted and unsorted cell populations. Genomic DNA from frozen cell pellets was isolated by Phenol/Chloroform extraction. Briefly, cell pellets were thawed and resuspended in 250 μl of Lysis buffer (1% SDS (Thermo Fisher Scientific), 0.2 M NaCl and 5 mM DTT (Roth) in TE Buffer) and incubated overnight at 65 °C. 200 μg of RNAse A (Thermo Fisher Scientific) were added to the sample and incubated at 37 °C for 1 h. 100 μg of Proteinase K (Sigma) were subsequently added followed by a 1 h incubation at 50 °C. Phenol/Chloroform/Isoamyl alcohol (Roth) was added to each sample in a 1:1 ratio, the mixture was vortexed for 1 min and subsequently centrifuged at 16,000 x g for 10 min at room temperature. The aqueous phase was transferred to a new tube, 1 ml 100% EtOH, 90 μl 5 M NaCl and 2 μl Pellet Paint (Merck) was added to each sample, mixed, and incubated at −80 °C for 1 h. DNA was pelleted by centrifugation for 16,000 x g for 15 min at 4 °C, pellets were washed twice with 70% EtOH, air-dried and resuspended in 50 μl H2O.

The genomically integrated sgRNA cassette was PCR-amplified to attach sequencing adaptors and sample barcodes. To ensure proper library coverage (300x), a total of 20 μg of each sample were amplified using the ReadyMix Kapa polymerase (Roche) with a total of 25 cycles and an annealing temperature of 56 °C. A relatively low amount of 0.5 μg genomic DNA was amplified per 50 μl PCR reaction since in samples stained with Flow-FISH, PCR amplification was inhibited at higher DNA concentrations. PCR was performed with the primer LR175 in combination with a sample-specific primer which contains a distinct 6-nucleotide barcode to allow sample identification after multiplexed deep sequencing (Primer sequences in Suppl. Table 4, LR178/LR180). Successful amplification was verified on a 1% agarose gel and the reactions were pooled. 1 ml of each pooled PCR was purified using the QIAquick PCR Purification Kit (Qiagen), loaded on a 1% agarose gel and purified using the QIAquick Gel Extraction Kit (Qiagen).

Libraries were sequenced as follows: replicate 1, paired-end 75 bp on the HiSeq 4000 platform, replicate 2, paired-end 50 bp on the HiSeq 2500 platform, replicate 3, single-read 75 bp on the HiSeq 2500 platform, using the custom primer LR176 yielding approximately 8*10^6^ fragments per sample (Read alignment statistics found in Suppl. Table 1).

#### Screen analysis

Data processing and statistical analysis was performed using the MAGeCK CRISPR screen analysis tools (Li et al., 2014, 2015) (v0.5.9.3). Alignment and read counting was performed with options [count --norm-method control] for samples from all 3 replicates together. At least 6.95*10^6^ mapped reads were obtained per sample. Correlation between the three replicates was computed as a Pearson correlation coefficient on the normalized counts (Suppl. Fig. 1D). The NTC distribution width was similar across samples, suggesting that sufficient library coverage was maintained during all steps (Suppl. Fig. 1E). Statistical analysis was performed in two steps. Since the CRISPRaX library often targets multiple TSSs per gene, with a subset of sgRNAs targeting multiple TSSs, we first identified one TSS per gene with the strongest effect. To this end, a first analysis was performed on the transcript level, including all TSS, with options [mle --norm-method control]. For each gene the TSS with the lowest Wald.fdr was identified. Then a statistical analysis was performed on the gene level, based on only those sgRNAs that targeted the identified TSS with options [mle --norm-method control]. Genes were ranked for their effect on *Xist* expression based on their beta score, a measure of the effect size estimated by the MAGeCK mle tool. For all visualization purposes the name Rnf12 was used for Rlim and Oct4 was used for Pou5f1. Alignment statistics, raw counts and gene hit summary files are provided in Suppl. Table 1.

### Bulk RNA-sequencing

Differentiating TX1072 XO mESCs (clone H7/A3) were profiled in three biological replicates by bulk RNA-seq as described previously for TX1072 XX mESCs (40). RNA-seq libraries were generated using the Tru-Seq Stranded Total RNA library preparation kit (Illumina) with 1 μg starting material for rRNA-depletion and amplified with 15 Cycles of PCR. Libraries were sequenced 2×50bp on a HiSeq 2500 with 1% PhiX spike-in, which generated ~50 Mio. fragments per sample.

### CUT&Tag

CUT&Tag experiments were performed on XEN and TS female cells as described previously (16). Cells were washed with PBS and dissociated with accutase. For each CUT&Tag reaction 1*10^5^ cells were collected and washed once with wash buffer (20 mM HEPES-KOH, pH 7.5, 150 mM NaCl, 0.5 mM spermidine, 10 mM sodium butyrate, 1 mM PMSF). 10 μl Concanavalin A beads (Bangs Laboratories) were equilibrated with 100 μl binding buffer (20 mM HEPES-KOH, pH 7.5, 10 mM KCl, 1 mM CaCl_2_, 1 mM MnCl_2_) and afterwards concentrated in 10 μl binding buffer. The cells were bound to the Concanavalin A beads by incubating for 10 min at room temperature with rotation. Following this, the beads were separated on a magnet and resuspended in 100 μl chilled antibody buffer (wash buffer with 0.05% digitonin and 2 mM EDTA). Subsequently 0.5 μl (GATA2/3/4/6 and IgG control) or 1 μl (H3K27ac, H3K27me3) of primary antibody was added and incubated on a rotator for 3 h at 4 °C. After magnetic separation the beads were resuspended in 100 μl chilled dig-wash buffer (wash buffer with 0.05% Digitonin) containing 1 μl of matching secondary antibody and were incubated for 1 h at 4 °C with rotation. The beads were washed three times with ice-cold dig-wash buffer and resuspended in chilled dig-300 buffer (20 mM HEPES-KOH, pH 7.5, 300 mM NaCl, 0.5 mM spermidine, 0.01% digitonin, 10 mM sodium butyrate, 1 mM PMSF) with 1:250 diluted 3xFLAG-pA-Tn5 preloaded with mosaic-end adapters. After incubation for 1 h at 4 °C with rotation, the beads were washed four times with chilled dig-300 buffer and resuspended in 50 μl tagmentation buffer (dig-300 buffer 10 mM MgCl_2_). Tagmentation was performed for 1 h at 37 °C and subsequently stopped by adding 2.25 μl 0.5 M EDTA, 2.75 ml 10% SDS and 0.5 μl 20 mg/mL Proteinase K and vortexing for 5 sec. DNA fragments were solubilized for 14 h at 55 °C followed by 30 min at 70 °C to inactivate residual Proteinase K. To remove the beads, the samples were put on a magnetic rack and the supernatants were transferred to a new tube. DNA fragments were purified with the ChIP DNA Clean & Concentrator kit (Zymo Research) and eluted with 25 μl elution buffer according to the manufacturer’s guidelines. Antibodies used can be found in Suppl. Table 4.

#### Library preparation and sequencing

NGS libraries were generated by amplifying 12 μl of the eluted CUT&Tag DNA fragments with i5 and i7 barcoded HPLC-grade primers (90) (Suppl. Table 4) with NEBNextHiFi 2x PCR Master Mix (New England BioLabs) on a thermocycler with the following program: 72 °C for 5 min, 98 °C for 30 s, 98 °C for 10 s, 63 °C for 10 s (14-15 Cycles for step 3-4) and 72 °C for 1 min. Post PCR cleanup was performed with Ampure XP beads (Beckman Coulter). For this 1.1x volume of Ampure XP beads were mixed with the NGS libraries and incubated at room temperature for 10 min. After magnetic separation, the beads were washed three times on the magnet with 80% ethanol and the libraries were eluted with Tris-HCl, pH 8.0. The quality of the purified NGS libraries was assessed with the BioAnalyzer High Sensitivity DNA system (Agilent Technologies). Sequencing libraries were pooled in equimolar ratios, cleaned again with 1.2x volume of Ampure XP beads and eluted in 20 μl Tris-HCl, pH 8.0. The sequencing library pool quality was assessed with the BioAnalyzer High Sensitivity DNA system (Agilent Technologies) and subjected to Illumina PE75 next generation sequencing on the NextSeq500 platform totalling 1-12 mio fragments per library (see Suppl. Table 3 for details).

### NGS data analysis

#### Published ChIP-seq data

FASTQ files for transcription factor binding data of FLAG-tagged GATA6 in mESCS after 36 hours of dox-mediated GATA6 overexpression (48) was retrieved from the GEO Accession Viewer (GSE69322) using *fasterq-dump* (v2.9.4) (http://ncbi.github.io/sra-tools/).

#### Data processing

For CUT&Tag and ChIP-seq data, reads were trimmed for adapter sequences using Trim Galore (0.6.4) with options [--paired --nextera] or [--paired --illumina] (http://www.bioinformatics.babraham.ac.uk/projects/trim_galore/) prior to alignment. Read alignment was performed to the mm10 reference genome using bowtie2 (v2.3.5.1) with options [--local --very-sensitive-local --no-mixed --no-discordant --phred33 -I 10 -X 2000] (Langmead and Salzberg, 2012) for CUT&Tag/ChIP-seq or with STAR (v2.7.5a) with options [--outSAMattributes NH HI NM MD] (Dobin et al., 2013) for RNA-seq. Sequencing data was then filtered for properly mapped reads and sorted using samtools (Li et al., 2009) (v1.10) with options [view -f 2 -q 20] (CUT&Tag/ChIPseq) or [view -q 7 -f 3] (RNA-seq) and [sort]. For ChIP-seq/CUT&Tag, blacklisted regions for mm10 (ENCODE Project Consortium, 2012) were removed using bedtools (Quinlan and Hall, 2010) (v2.29.2) with options [intersect -v]. For ChIP-seq, reads were also deduplicated using Picard (v2.18.25) with options [MarkDuplicates VALIDATION_STRINGENCY=LENIENT REMOVE_DUPLICATES=TRUE] (http://broadinstitute.github.io/picard).

Mapping statistics and quality control metrics for RNA-seq/CUT&Tag can be found in Suppl. tables 2 and 3.

#### Generation of coverage tracks

BIGWIG coverage tracks for CUT&Tag and ChIP-seq were created using deeptools2 (v3.4.1) (Ramírez et al., 2016) on merged replicates with the options [bamCoverage -bs 10 -e --normalizeUsing CPM -ignore chrX chrY]. The tracks were visualized using the UCSC genome browser (Kent et al., 2002).

#### Peak calling

Peaks for CUT&Tag and ChIP-seq were called using MACS2 (Zhang et al., 2008) (v2.1.2) with standard options [callpeak -f BAMPE/BAM -g mm -q 0.05] on individual replicates. For ChIP-seq, input samples were included for normalization using [-c]. Only peaks detected in all replicates were retained by merging replicates using bedtools (Quinlan and Hall, 2010) (v2.29.2) with [intersect].

#### Correlation analysis

For CUT&Tag, BAM files, excluding mitochondrial reads, were counted in 1 kb bins using deepTools2 (Ramírez et al., 2016) (v3.4.1) with options [multiBamSummary bins -bs 1000 -bl chrM.bed]. The Pearson correlation coefficient between different histone marks, conditions or replicates was then computed using deepTools2 (v3.4.1) with options [plotCorrelation -c pearson]. The resulting values were hierarchically clustered and plotted as a heatmap. Correlation between biological replicates was high and the samples showed the expected correlation patterns (Suppl. Fig. 5B).

#### Annotation of GATA factor motifs within CUT&Tag peaks within the Xic

To identify peaks detected by CUT&Tag for GATA TFs that contain the respective GATA binding motif, FASTA files containing the sequences of all peaks that were identified in both replicates were generated using bedtools (v2.29.2) (91) with options [getfasta]. The FASTA files were scanned for the occurence of the respective GATA transcription factor binding motif, which were retrieved from the JASPAR database (92) (8^th^ release) using FIMO with options [--thresh 0.001] (93). The location and annotation of all peaks within the *Xic* is shown in Supplemental Table 3.

#### Verification of GATA CUT&Tag data

As GATA factors have, to our knowledge, not been profiled by CUT&Tag previously, we first assessed the data quality. To assess specificity of the identified peaks, we compared the intensity of peaks with a GATA motif to those without. To this end, we used RSubread (Liao et al., 2019) (v2.0.1) with options [featureCounts(isPairedEnd = TRUE)] to count the number of reads mapping to peaks with or without a motif individually. Subsequently, we transformed these counts into Reads per Million (RPM) and plotted their density (Suppl. Fig. 5C). While peaks with a motif were clearly stronger for GATA6, and to a slightly lesser extent also for GATA3 and GATA4, no difference was observed for GATA2 (Suppl. Fig. 5C).

As another quality control step we identified enriched motifs within all peaks of each CUT&Tag data set. To this end, we performed a motif enrichment using the non-redundant vertebrate JASPAR2020 CORE position frequency matrix (PFM) dataset, as described previously (94) with adaptations, to see if we could recover GATA-factor associated motifs. To this end, all peaks that were identified in both replicates were centered and extended to a total of 500 bp. Afterwards, Rsubread (Liao et al., 2019) (v2.0.1) with options [featureCounts(isPairedEnd = TRUE)] was used to quantify the number of reads mapping to each peak. The centered peaks were ranked depending on RPM and transformed into FASTA files using bedtools (v2.29.2) (91) with options [getfasta]. These files were scanned for enriched PFMs using AME (McLeay & Bailey 2010) with options [--shuffle]. For GATA3, GATA4 and GATA6 all top-ranking motifs were members of the GATA family, while no GATA motifs were found for GATA2. These analyses suggest that GATA3, GATA4 and GATA6 can be profiled reliably by CUT&Tag, while the data for GATA2 should be interpreted with caution. The complete results of the motif enrichment analysis are shown in Suppl. Table 3.

#### Gene quantification of RNA-seq data

RNA-seq data during the differentiation of female TX1072 mESCs (XX) was acquired from GSE151009 (40). (Single-cell)-RNA-seq data during mouse embryonic development (Deng et al. 2014; Zhang et al. 2018) was similarly acquired from GEO (GSE45719, GSE76505). The single-cell data (Deng et al. 2014) was merged as a pseudo-bulk prior to alignment. Gene expression was quantified using the GENCODE M25 annotation (Frankish et al., 2019). Rsubread (Liao et al., 2019) (v2.0.1) was used with the options [featureCounts(isPairedEnd = TRUE, GTF.featureType = “exon”, strandSpecific = 2)]. TPM values for the XX and XO time courses can be found in Supplementary Table 3.

### Single-cell RNA-seq analysis

For reanalysis of previously published scRNA-seq data from mouse embryos, the normalized data from study of preimplantation embryos up to E3.5 (41) was downloaded from GEO (GSE45719) and data from E4.5-E6.5 embryos (56) was downloaded from https://github.com/rargelaguet/scnmt_gastrulation together with the cell type annotation and visualized in R.

### Data availability

CRISPRa screen, CUT&Tag and TX1072 XO bulk RNA-seq data sets are available via GEO (GSE194018). The scRNA-seq data from mouse embryos analyzed in this study (41,56), are available on Github (https://github.com/rargelaguet/scnmt_gastrulation) and on GEO (GSE121708, GSE45719). Bulk RNA-seq for TX1072 XX (40) and ChIP-seq for FLAG-tagged GATA6 (48) are available via GEO (GSE151009, GSE69323).

## Acknowledgements

We are thankful for the support and feedback received from members of the Schulz lab. We want to thank Pablo Navarro for sharing the E14 SunTag cell line (E14-STN) and Joost Gribnau and Catherine Dupont for sharing the female XEN cell line. We thank Maud Borensztein for support in setting up RNA-FISH on mouse preimplantation embryos and Edith Heard and Christel Picard for sharing genomic DNA of IKE15-9TG80 and IKE14-2TG53 mESCs. We thank Vera Schmiedel for cloning RE-targeting multiguide plasmids and Verena Mutzel and Benedikt Boesen for cloning SP265. We thank the Max Planck Institute for Molecular Genetics Seqcore, Flow Cytometry, Imaging, Transgene and animal facilities. Specifically, we thank Judith Fiedler, Mirjam Peetz, Adrian Landsberger, Christin Franke, and members of the transgenic animal facility. This work was supported by the Max-Planck Research Group Leader program, E:bio Module III—Xnet grant (BMBF 031L0072) and Human Frontiers Science Program (CDA-00064/2018) to E.G.S. T.S. was supported by the DFG (IRTG2403, Regulatory Genome) and G.N. by the European Union’s Horizon 2020 Research and Innovation Program (Marie Skłodowska-Curie ITN PEP-NET).

## Author Contributions

L.R.L. and E.G.S. conceived the project and designed the experiments. L.R.L. performed most experiments with help from I.D., A.G. and G.N. *Gata1/4/6* TKO in embryos was performed by A.S.K. and L.W., IF staining of embryos with help from M.S. and RNA-FISH in embryos by L.R.L and I.D. T.S. analyzed CRISPRa screen, bulk RNA-seq, CUT&Tag and GATA6 ChIP-seq. I.D. performed CUT&Tag, immunofluorescence-RNA-FISH and NGS karyotyped all cell lines. R.W. generated TSC RE79 heterozygous KO lines and maintained TSC cultures. G.P. designed the CRISPRaX sgRNA library. E.G.S. analyzed scRNA-seq data. G.N. performed and analyzed the enhancer reporter assay. R.B. and E.G.S. analyzed images and data from immonofluorescence-RNA-FISH and RNA-FISH on embryos. L.R.L, T.S., and E.G.S. wrote the manuscript with input from all authors. Funding acquisition by A.M. and E.G.S.

## Supplemental Figures

**Suppl. Fig 1:**
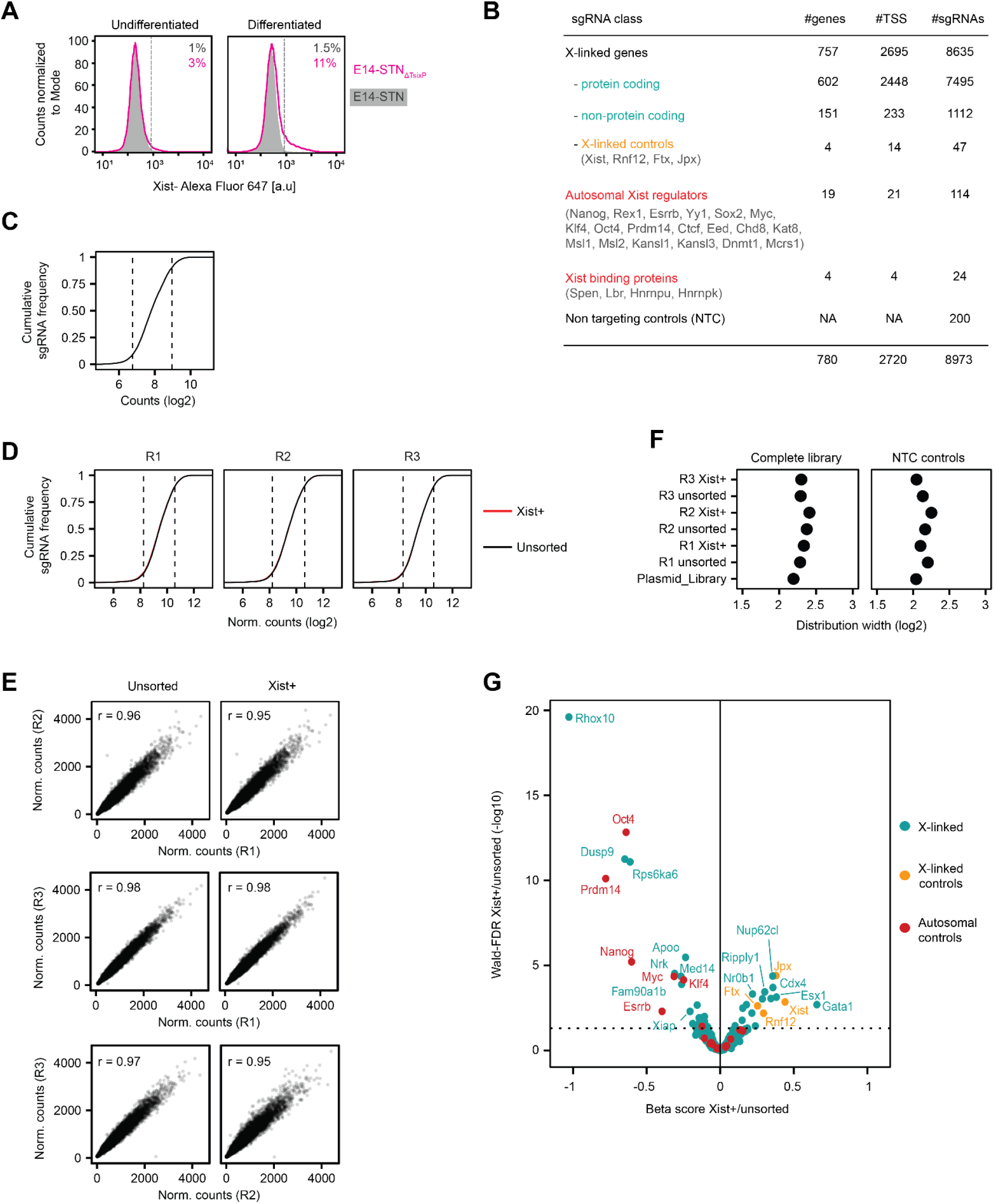
Pooled CRISPR activation screen identifies new Xist regulators. **(A)** E14-STN (grey) and E14-STN_ΔTsixP_ (pink) cells were treated with doxycycline for 3 days and were differentiated for the last 2 days by LIF withdrawal, followed by Flow-FISH with Xist-specific probes. Deletion of *Tsix* major promoter did not lead to *Xist* ectopic expression upon differentiation. Dashed lines mark the 99th percentile of undifferentiated E14-STN cells to separate Xist+ and Xist-cells. The percentage of Xist+ cells in each sample is indicated. **(B)** Composition of the CRISPRaX sgRNA library, targeting each TSS with 6 sgRNAs per gene. Since a subset of guides target different coding and non-coding transcripts, the total number of sgRNAs is smaller than the sum of sgRNAs across categories. **(C-D)** Cumulative frequency plot showing the distribution of sgRNA counts in the cloned sgRNA library (C) and in the sorted and unsorted fractions (D). Dashed lines indicate the distribution width (10th and 90th percentile, quantified in F). **(E)** Scatterplots showing a high correlation between the replicates in the screen for each fraction as indicated. Pearson correlation coefficients between replicates are shown. **(F)** Log2 distribution width (fold change between the 10th and 90th percentiles) for all sgRNAs (left) and non-targeting (NTC) sgRNAs only (right). The NTC distribution width was similar across samples, suggesting that sufficient library coverage was maintained during all steps of the screen. **(G)** Volcano plot of the screen results, showing the beta-score as a measure of effect size vs Wald-FDR (MAGeCK-MLE), colored according to gene class as indicated. The dotted line denotes Wald-FDR <0.05

**Suppl. Fig 2:**
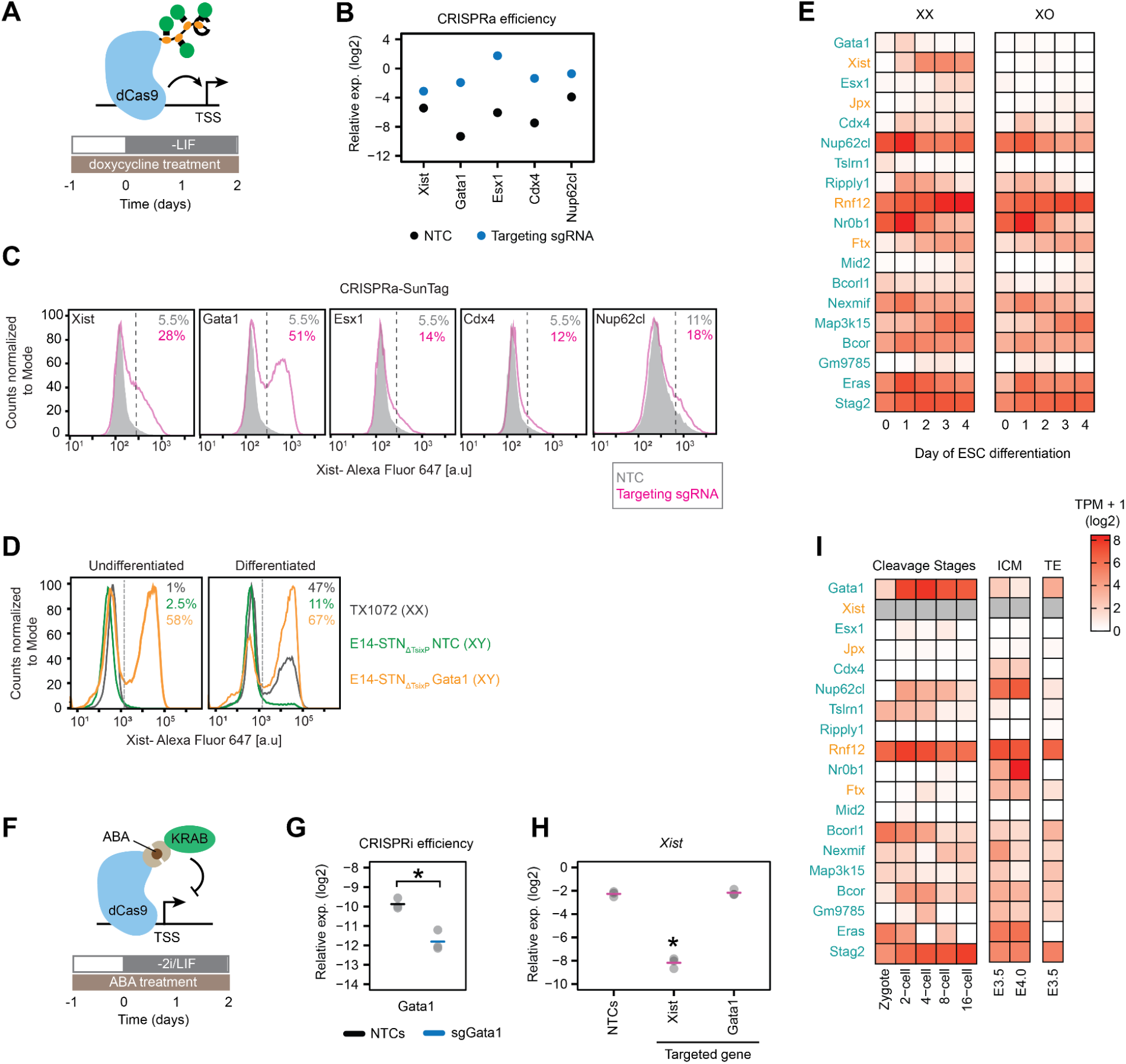
GATA1 is a potent *Xist* activator. **(A-C)** Individual overexpression of screen hits with CRISPRa in E14-STN_ΔTsixP_ mESCs using a single guide RNA per gene that had performed well in the screen. **(A)** The cells were treated with doxycycline 24 h before differentiation by LIF withdrawal for 2 days. **(B)** Expression levels of the targeted genes were measured by qRT-PCR. **(C)** *Xist* expression measured by Flow-FISH. Dashed lines mark the 99th percentile of undifferentiated NTC-transduced E14-STN_ΔTsixP_ cells (Xist-population). The percentage of Xist+ cells is indicated. **(D)** *Xist* expression was measured via Flow-FISH in female TX1072 cells line and in E14-STN_ΔTsixP_ cells transduced with multiguide expression vectors of three sgRNAs against *Gata1* promoter region or with non-targeting controls (NTC). TX1072 cells were cultured in naive conditions (2i/LIF) and E14-STN_ΔTsixP_ in conventional ESC medium (LIF). The cells were differentiated (2i/LIF or LIF withdrawal) for 2 days. E14-STN_ΔTsixP_ were treated with doxycycline 24 h before and during differentiation. Dashed lines mark the 99th percentile of the TX1072 undifferentiated (2i/LIF) sample and the percentage of Xist+ cells in each sample is indicated. **(E)** Heatmap showing expression levels assessed by RNA-seq (mean of 3 biological replicates) of the most enriched genes in the screen (Fig. 1B) in XX and XO TX1072 mESCs differentiated by 2i/LIF withdrawal. **(F-H)** *Gata1* knock-down by CRISPRi in female mESCs. (F) Schematic representation of an ABA-inducible CRISPRi system in female TX-SP107 mESCs. **(G-H)** *Gata1* knock-down (G) and effect on *Xist* (H) quantified by qRT-PCR after 2 days of differentiation. SgRNAs targeting the *Xist* TSS and NTCs were included as controls. Horizontal dashes indicate the mean of 3 biological replicates (dots); asterisks indicate p<0.05 for paired Student’s T-test. **(I)** Expression of screen hits during preimplantation development (41,42). *Xist* could not be quantified (grey) because the employed protocol was not strand-specific, such that *Xist* could not be distinguished from its antisense transcript *Tsix*. In (E) and (I) *Xist* and known Xist regulators are colored in yellow.

**Suppl. Fig 3:**
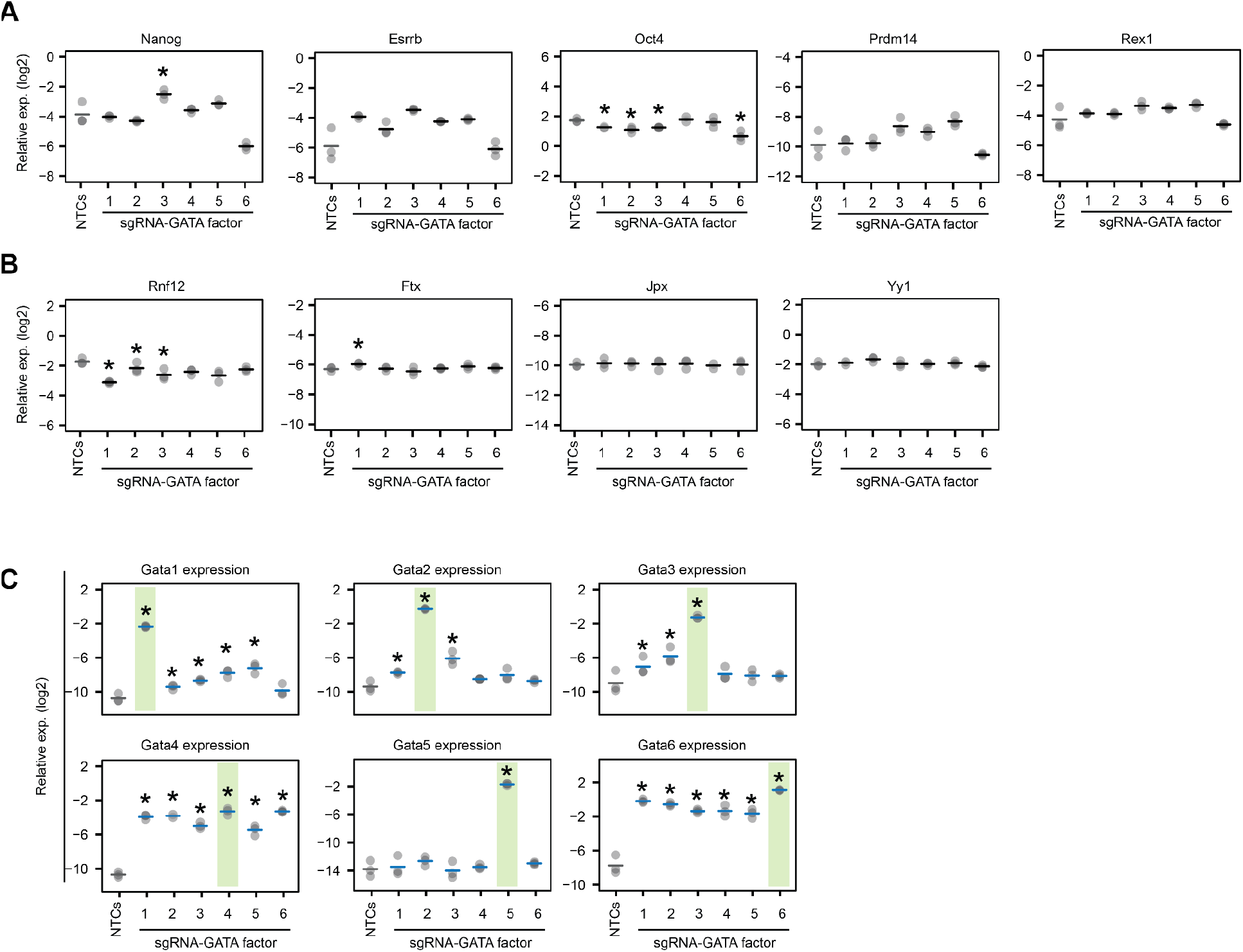
CRISPRa-mediated overexpression of GATA TFs. **(A-C)** Male E14-STN_ΔTsixP_ cells were transduced with multiguide expression vectors of three sgRNAs targeting the promoter of each GATA factor or with non-targeting controls (NTCs). Cells were treated with doxycycline for 3 days and differentiated for 2 days (LIF withdrawal). Expression levels of pluripotency factors (A) known Xist regulators (B) and of GATA factors (C) were assessed by qRT-PCR. Mean (horizontal dashes) of 3 biological replicates (dots) is shown; asterisks indicate p<0.05 of a paired Student’s T-test for comparison to the respective NTC control. Green areas in (C) indicate the GATA factor that was targeted by CRISPRa.

**Suppl. Fig. 4:**
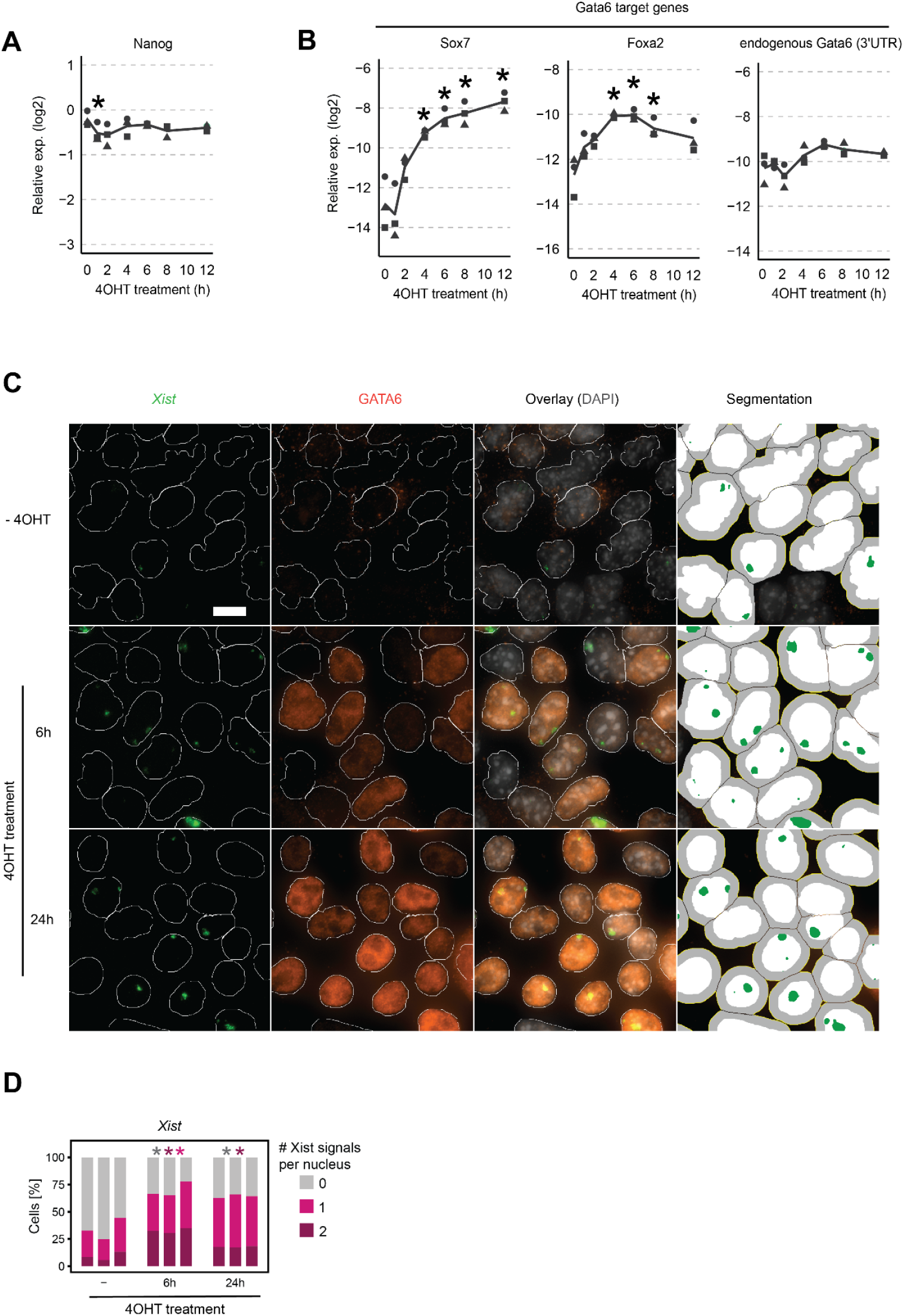
*Xist* is rapidly induced by GATA6 in a dose-dependent manner. **(A-B)** Time course of 4OHT treatment of TX-SP107 ERT2-Gata6-HA cells, cultured in 2i/LIF medium. Expression levels of Nanog (A) or known GATA6 target genes (B) were measured via qRT-PCR. The line indicates the mean of 3 biological replicates (symbols); asterisks indicate p<0.05 using a paired Student’s T-test, comparing levels to the untreated control (0h). **(C-D)** TX-SP107 ERT2-Gata6-HA cells, cultured in 2i/LIF medium were grown on glass coverslips in conventional ESC medium (LIF only) for 48h and treated with 4OHT for 6 or 24h, followed by immunofluorescence staining (anti-HA to detect GATA6) combined with RNA-FISH (to detect *Xist*). Since cells cultured in 2i/LIF grow in tight colonies, cells had to be cultured without 2i for 48h, where they flatten out and thus facilitate the analysis. Nuclei (white) and *Xist* signals (green) were detected by automated image segmentation and GATA6-HA staining was quantified in the nucleus and in a 2.64 μm ring around the nucleus as proxy for the cytoplasm (right column, grey). (D) The number of *Xist* signals per nucleus is shown. 2i removal led to partial *Xist* derepression, such that 25-44% of cells already expressed *Xist* without 4OHT treatment, which increased to 65-78% after 6h OHT treatment. While about half of the *Xist*-positive cells expressed *Xist* from both alleles after 6h, the majority of cells showed a mono-allelic pattern after 24 h. Such a biallelic-to-monoallelic transition, probably mediated by negative feedback regulation, is also observed in differentiating ES cells (40) and is in agreement with previous reports of random monoallelic *Xist* upregulation, when ES cells are differentiated into XEN cells by GATA6 overexpression (95). 3 biological replicates are shown, with excluding nuclei with >2 Xist signals. Asterisks indicate p<0.05, paired Student’s T-test compared to −4OHT sample. Scale bar represents 10 μm.

**Suppl. Fig 5:**
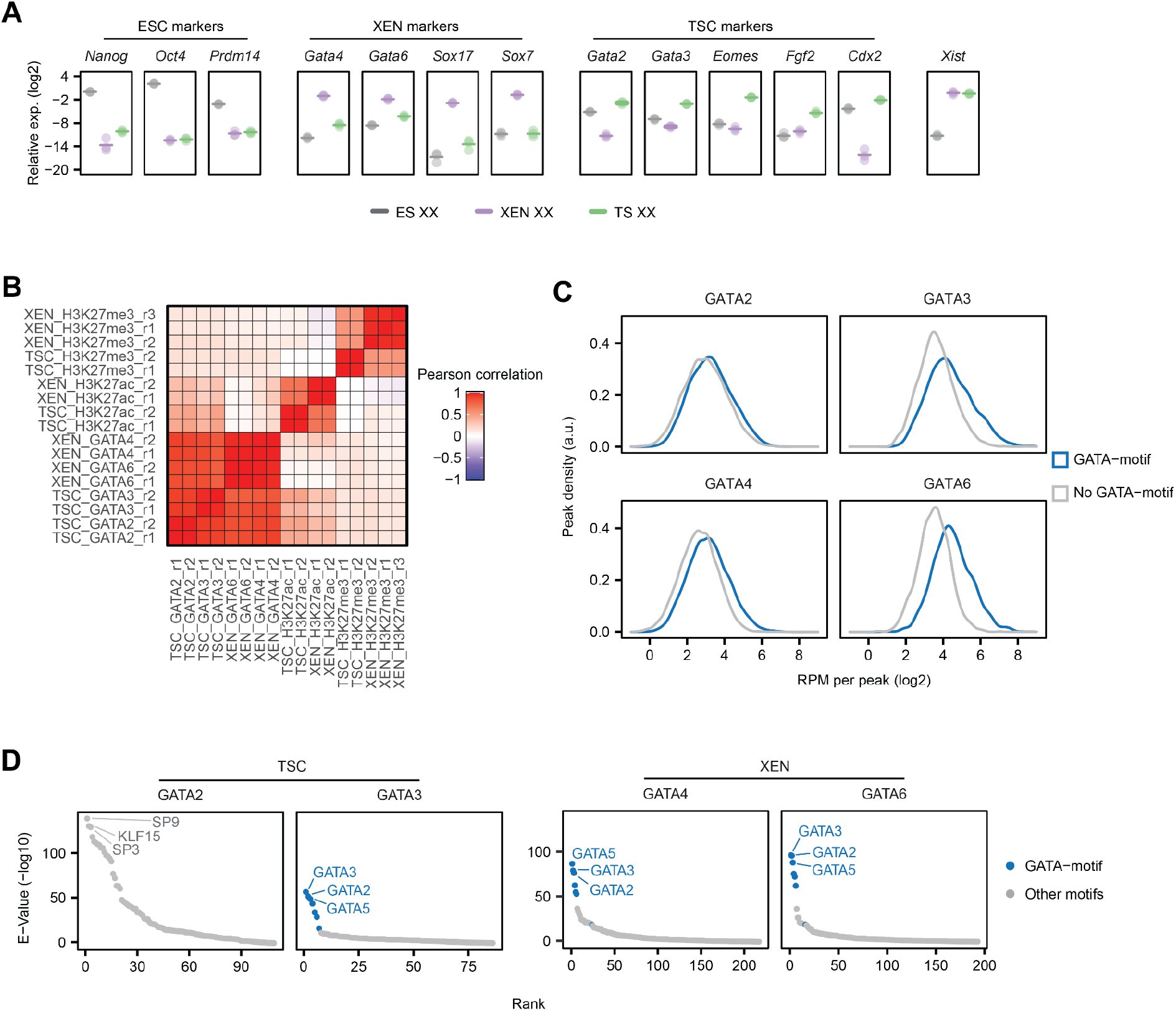
GATA factor profiling by CUT&Tag in XEN and TSCs. **(A)** Relative expression levels of various marker genes of ESCs, XEN and TS cells as indicated and of *Xist*, measured via qRT-PCR in female TX1072 ESCs, XEN and TS cells. Mean (dash) of 3 biological replicates (dots) is shown. **(B)** Pearson correlation coefficient between all CUT&Tag samples. The heatmap is ordered according to hierarchical clustering of the correlations. Correlation between biological replicates was high and the samples showed the expected correlation patterns. **(C)** Density of RPM values per peak in each condition of the GATA CUT&Tag. The data is split in peaks containing (blue) or not containing (grey) the respective GATA-motif (p < 0.001, FIMO). While peaks with a motif were clearly stronger for GATA6 and GATA3, and to a slightly lesser extent also for GATA4, no difference was observed for GATA2. **(D)** Enrichment of TF-binding motifs within peaks identified for the different GATA TFs using AME. Binding motifs were ranked according to their E-values, a measure of the statistical enrichment of the respective motif. All binding motifs with an −log10(E-value) < 10 are shown. All GATA-family binding motifs are colored in blue. Additionally, the 3 most enriched motifs per sample are labeled. For GATA3, GATA4 and GATA6 all top-ranking motifs were members of the GATA family, while no GATA motifs were found for GATA2. These analyses suggest that GATA3, GATA4 and GATA6 can be profiled reliably by CUT&Tag, while the data for GATA2 should be interpreted with caution.

**Suppl. Fig 6:**
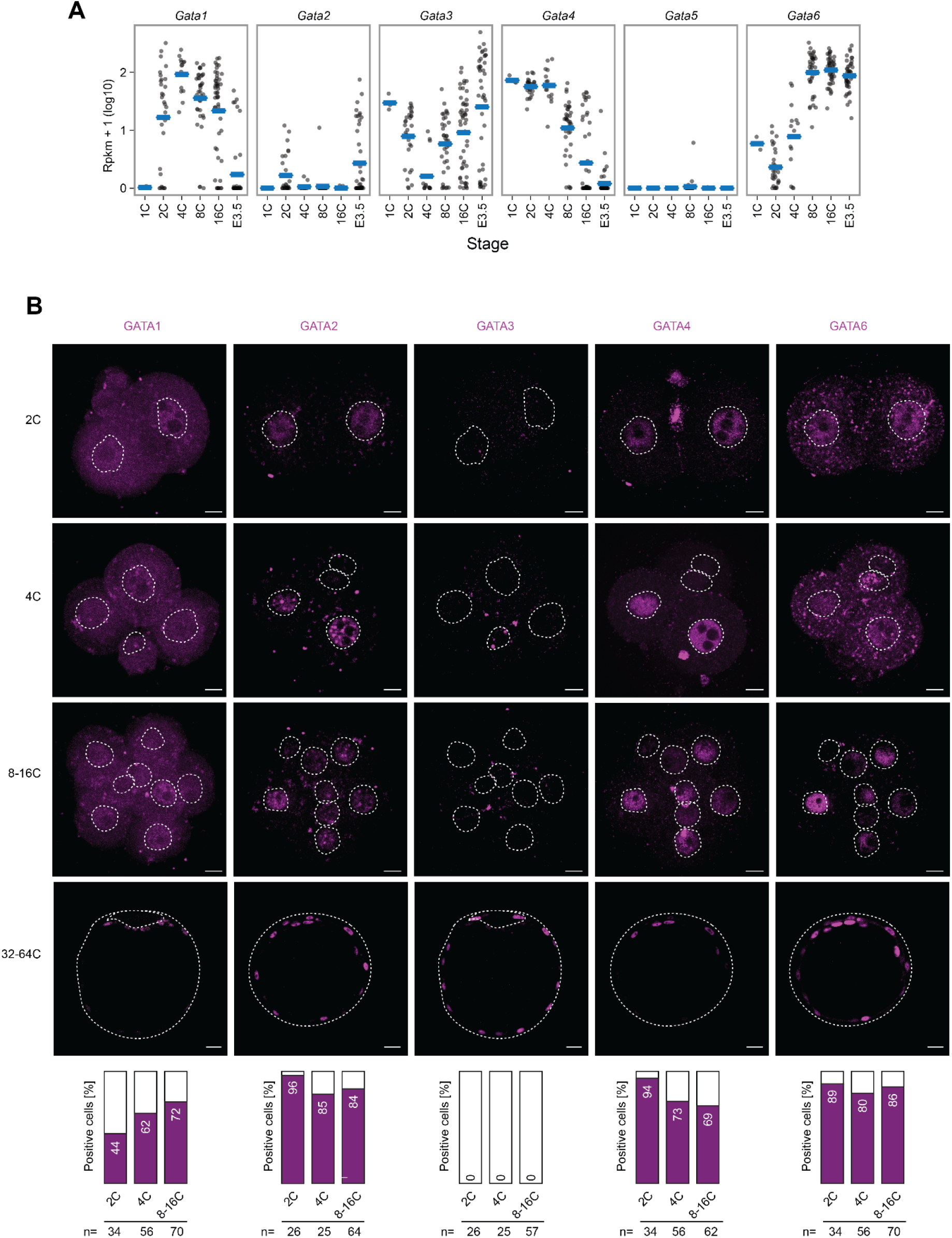
Multiple GATA TFs are expressed during mouse preimplantation development. **(A)** Expression of GATA TFs assessed by scRNA-seq across different stages of early mouse development (41). Horizontal dashes indicate the mean of 24 (1C), 180 (2C), 84 (4C), 222 (8C), 300 (16C) and 258 (E3.5) cells. **(B)** Protein staining of all GATA TFs except GATA5 in preimplantation mouse embryos (stages indicated). Nuclei were detected by DAPI staining and their contour is marked (dashed line). Bar plots show the percentages of positive nuclei for the respective GATA protein. Percentages represent the mean of two biological replicates. The number of nuclei counted is shown below the plots. Scale bars represent 10 μm, scale bars for 32-64C are 20 μm.

**Suppl. Fig. 7.**
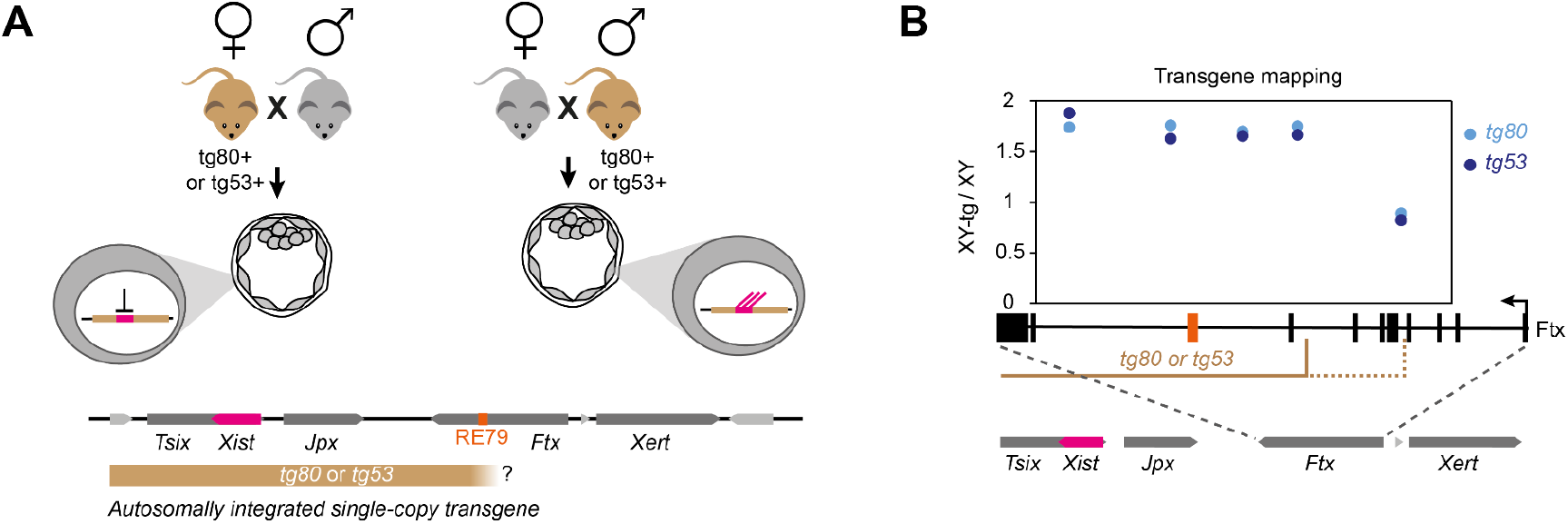
RE79 is part of the tg80/tg53 transgenes, which can drive post-fertilization *Xist* upregulation. **(A)** The *tg80/tg53* transgenes (beige), which contain the *Xist* gene and ~100 kb of upstream genomic sequence (bottom), can reproduce imprinted *Xist* expression, when autosomally integrated as a single copy, as they are expressed upon paternal (right), but not upon maternal (left) transmission (17,18). **(B)** Mapping of the telomeric end of *tg80/tg53* by qPCR on genomic DNA from XY-tg80/tg53 ESCs with primer pairs detecting different positions around RE79, as indicated below the plot. Mapping confirms that *tg80* and *tg53* contain the RE79 region. Results are expressed as relative DNA quantity with respect to XY cells without the transgene (E14-STN_ΔTsixP_).

**Suppl. Fig 8:**
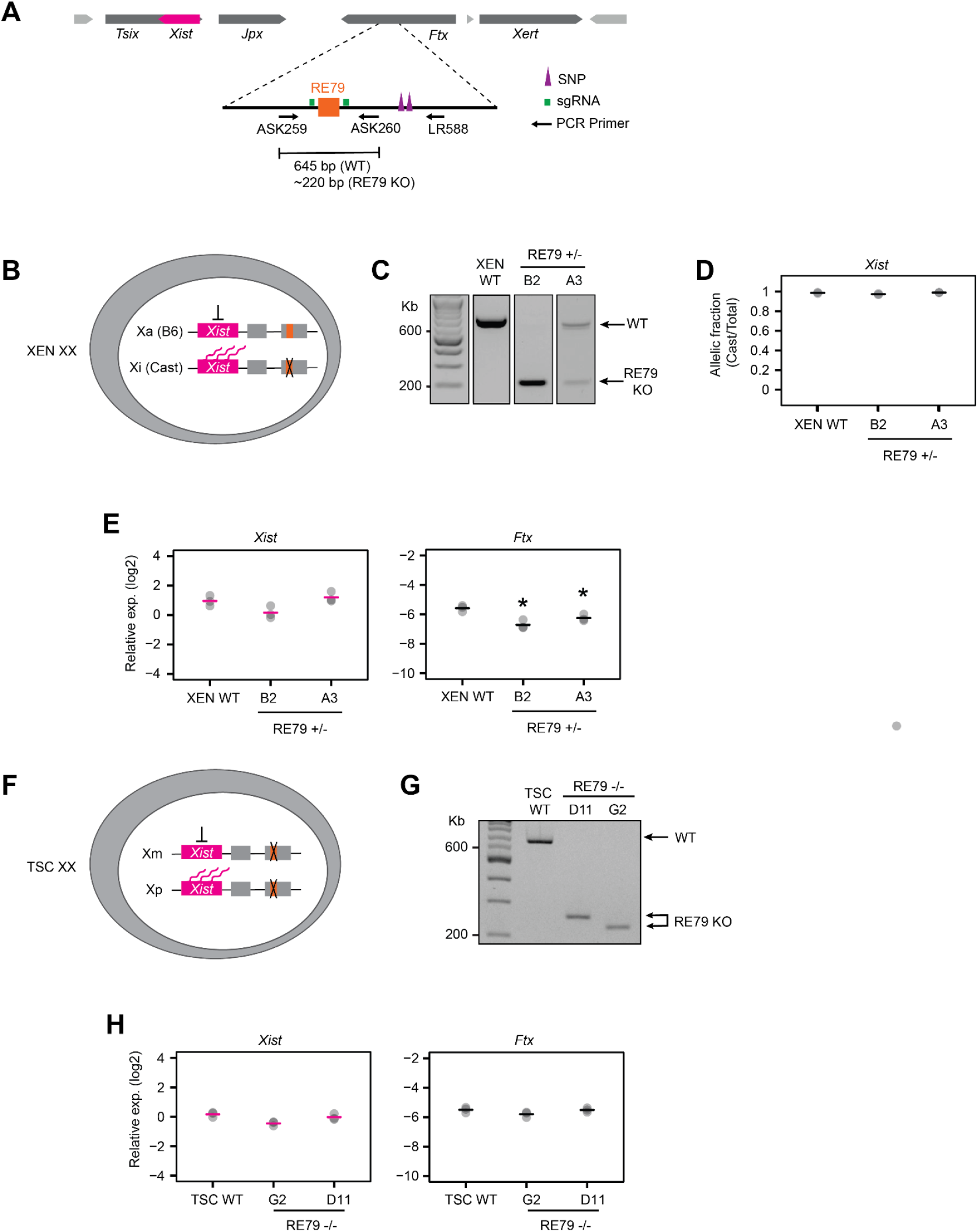
Deletion of RE79 in XEN and TS cells does not affect *Xist* expression. **(A)** Schematic representation of the genomic region surrounding RE79 (orange). 2 sgRNAs (green lines) were used to delete RE79 via the CRISPR/Cas9 system. Primers (ASK259/ASK260) used to genotype the clones are indicated. PCR that covers SNPs (purple triangles, primers ASK259/LR588) was employed to determine the deleted allele in XEN cells. **(B-E)** Generation of RE79 heterozygous deletion in female hybrid (B6/Cast) XEN cell line. Two RE79 knock-out clones (B2, A3) in which the paternal (Cast) locus was deleted, were analysed. **(C)** Agarose gel image of genotyping PCR. Sequencing of a PCR product that covers SNPs determined that RE79 was only deleted on the Cast allele in the B2 clone. **(D)** Pyrosequencing analysis on XEN RE79+/- clones confirming that *Xist* is expressed from the paternal X (Cast) chromosome. **(E)** qRT-PCR analysis of *Xist* and *Ftx* in XEN RE79+/- clones. Asterisks indicate p<0.05 of a paired Student’s T-test comparing each sample to the wildtype control. No effect on *Xist* expression could be detected in the clones, where RE79 had been deleted. **(F-H)** Generation of RE79 homozygous deletion in a female TS cell line. Genotyping PCR confirmed the deletion **(G)** and qRT-PCR was employed to measure *Xist* and *Ftx* expression levels **(H)**, showing no change in *Xist* expression upon RE79 deletion. In (D,E,H) individual measurements of 3 biological replicates are shown (dots) and the mean is indicated by a bar.

**Suppl. Fig 9:**
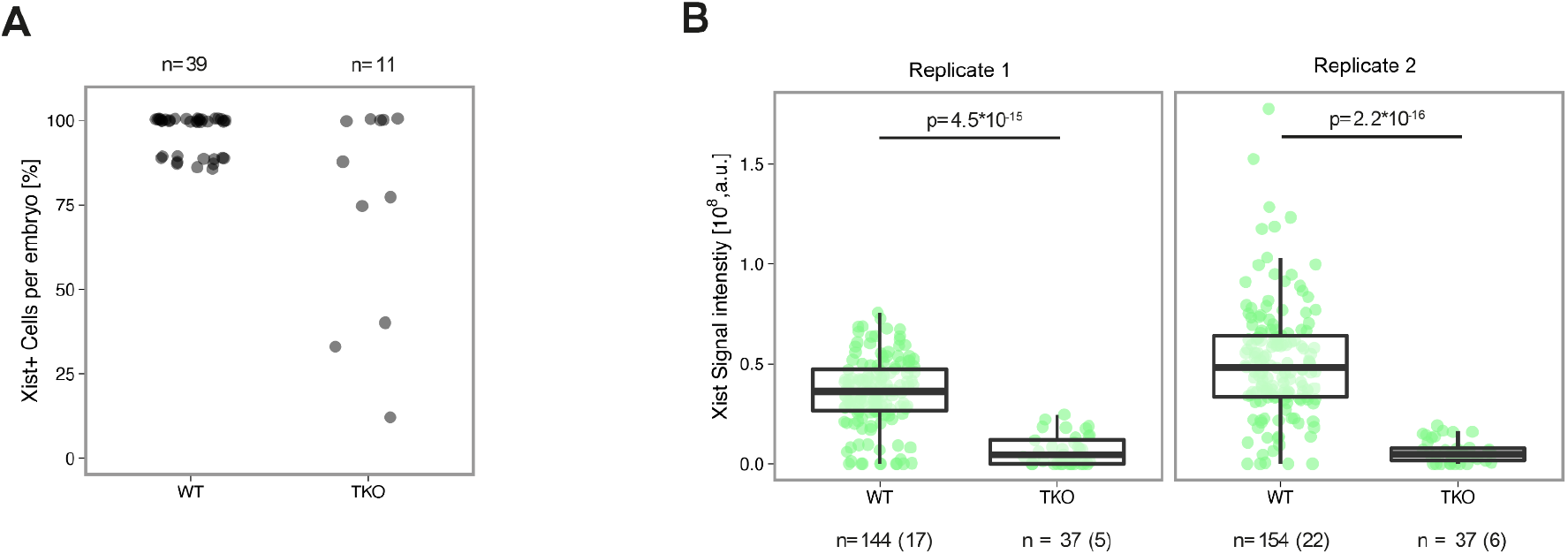
GATA1/4/6-TKO embryos exhibit inpaired *Xist* upregulation. Zygotic triple knock-out (TKO) of *Gata1, Gata4* and *Gata6* as shown in main Fig. 5 C. **(A)** The percentage of cells in each embryo with an *Xist* signal is shown at the 8-cell stage. Two biological replicates were merged. The efficiency of *Xist* upregulation is reduced in TKO embryos. **(B)** The summed fluorescence intensity within the automatically detected *Xist* cloud is shown for individual cells. Statistical comparison was performed with a Wilcoxon ranksum test. The number of cells (embryos) included in the analysis is indicated on top.

## Supplementary Tables

**Supplementary Table 1**. CRISPRaX sgRNA library and screen analysis. Related to Fig. 1 and Suppl. Fig. 1.

**Supplementary Table 2**. Bulk RNA-seq on TX1072 XX and XO cells. Related to Suppl. Fig 2..

**Supplementary Table 3**. CUT&Tag analysis and AME motif discovery results. Related to Fig. 4 and Suppl. Fig. 5.

**Supplementary Table 4**. Cell lines, oligos, probes, antibodies used in this study

**Supplementary Table 5.** Sequences of enhancer reporter plasmids and the ERT2-GATA6-HA expression vector.

